# Selective Requirement for Polycomb Repressor Complex 2 in the Generation of Specific Hypothalamic Neuronal Sub-types

**DOI:** 10.1101/2021.07.28.454060

**Authors:** Behzad Yaghmaeian Salmani, Brad Balderson, Susanne Bauer, Helen Ekman, Annika Starkenberg, Thomas Perlmann, Michael Piper, Mikael Bodén, Stefan Thor

## Abstract

The hypothalamus displays staggering cellular diversity, chiefly established during embryogenesis by the interplay of several signalling pathways and a battery of transcription factors. However, the contribution of epigenetic cues to hypothalamus development remains unclear. We mutated the Polycomb Repressor Complex 2 gene *Eed* in the developing mouse hypothalamus, which resulted in the loss of H3K27me3; a fundamental epigenetic repressor mark. This triggered ectopic expression of posteriorly expressed regulators (e.g., Hox homeotic genes), upregulation of cell cycle inhibitors and reduced proliferation. Surprisingly, despite these effects, single cell transcriptomic analysis revealed that the majority of neuronal subtypes were still generated in *Eed* mutants. However, we observed an increase in Glutamatergic/GABAergic double-positive cells, as well as loss/reduction of dopamine, Hypocretin/Orexin and Tac2 neurons. These findings indicate that many aspects of the hypothalamic gene regulatory flow can proceed without the key H3K27me3 epigenetic repressor mark, and points to a unique sensitivity of particular neuronal sub-types to a disrupted epigenomic landscape.

## INTRODUCTION

The hypothalamus is a small brain structure, which, in spite of its size acts as a master homeostatic regulator; controlling energy and fluid balance, thermoregulation, sleep-wake states, stress responses, growth and reproduction, as well as emotional and social behaviours (Saper and Lowell, 2014). The hypothalamus can perform this plethora of complex functions in large part due to its staggering neuronal diversity (Alpar et al., 2019; Romanov et al., 2019). Microarray and single cell transcriptomic analyses of the adult hypothalamus have resulted in major leaps in our understanding of the complete mature cellular diversity in this tissue, pointing to the presence of ∼50 major cell types and several hundred subtypes (Campbell et al., 2017; Chen et al., 2017; Dalal et al., 2013; Henry et al., 2015; Jeong et al., 2016; Kim et al., 2019; Kurrasch et al., 2007; Lam et al., 2017; Mickelsen et al., 2019; Mickelsen et al., 2017; Moffitt et al., 2018; Romanov et al., 2017; Shimogori et al., 2010).

The development of the hypothalamus has been challenging to decode, not only due to its immense cellular diversity. Unlike other CNS regions, such as the cortex, hindbrain or spinal cord, which are arranged in columnar structures, the adult hypothalamus is characterized by a patchwork of partially overlapping nuclei and territories. Hypothalamic development is also characterized by complex tissue rearrangements and cellular migration (Bedont et al., 2015; Burbridge et al., 2016; Ferran et al., 2015; Puelles and Rubenstein, 2015). Despite these challenges, extensive efforts have resulted in the unravelling of a multi-step process of anterior-posterior and medio-lateral patterning events, involving many of the major signalling pathways (Shh, BMP, Nodal, WNT, FGF). This patterning process results in, and integrates with, the selective expression of a number of early transcription factors (TFs), which act to further sub-divide the hypothalamus. These early TFs in turn activate panels of late TFs within subdomains of the developing hypothalamus, which act with more restrictive mandates to specify diverse subsets of hypothalamic cell fates (Alvarez-Bolado, 2019; Bedont et al., 2015; Blackshaw et al., 2010; Burbridge et al., 2016; Ferran et al., 2015; Nesan and Kurrasch, 2016; Puelles and Rubenstein, 2015; Xie and Dorsky, 2017). More recently, single cell transcriptomic analysis of the embryonic hypothalamus has greatly increased our understanding of its developmental process (Huisman et al., 2019; Kim et al., 2020; Romanov et al., 2020; Zhang et al., 2021; Zhou et al., 2020).

In contrast to the identification of key signalling cues and TF pathways, the role of epigenetics in the control of hypothalamus development is not well understood. An intensively studied epigenetic regulator is the Polycomb Group complex (PcG), which is a collective name that refers to several different sub-complexes, where the Polycomb Repressor Complexes 1 and 2 (PRC1 and PRC2) are arguably most well-defined (Piunti and Shilatifard, 2016; Steffen and Ringrose, 2014). PRC2 mono-, di-and tri-methylates residue K27, chiefly on Histone 3.3, and to a lesser extent H3.2 and H3.1 (Banaszynski et al., 2013). In general, PRC2 triggers transcriptional repression of target genes, although its specific role in gene regulation and chromatin compaction is still under intensive investigation, an issue that is further challenged by the existence of variations in PRC2 protein complex composition (Chammas et al., 2020; Laugesen et al., 2019; van Mierlo et al., 2019). In mammals, the *Embryonic ectoderm development* (*Eed*) gene constitutes a critical component of PRC2 and is encoded by a single gene in the mouse genome. *Eed* null mutants display an apparently complete loss of H3K27me1/2/3 (Montgomery et al., 2005), and when *Eed* is selectively removed in e.g., the hematopoietic lineage, in the intestine or in the CNS, H3K27me3 is lost (Jadhav et al., 2020; Xie et al., 2014; Yaghmaeian Salmani, 2018). Hence, in contrast to most, if not all other epigenetic marks and their enzyme systems, the single gene removal of *Eed* provides an unparalleled means of completely removing this key epigenetic mark.

To begin addressing the role of epigenetic input on hypothalamic development, we analysed *Eed* conditional mutants (*Eed-cKO*), where *Eed* was deleted by the early CNS deleter *Sox1-Cre*, which is active at E8.5 (Takashima et al., 2007). We find that *Eed-cKO* mutants display a complete loss of H3K27me3 in the hypothalamus, from E11.5 and onwards. *Eed-cKO* mutants upregulate several cell cycle inhibitor genes and display reduced proliferation in the hypothalamus. We also observed ectopic expression of many posteriorly expressed TFs, indicating posteriorization of the anterior CNS. To unravel the effects of *Eed-cKO* upon cell specification, we conducted single cell transcriptomic (scRNA-seq) analysis at E13.5, E15.5 and E18.5, spanning the major phase of hypothalamic cell specification (Kim et al., 2020; Romanov et al., 2020). Surprisingly, in spite of reduced proliferation and extensive ectopic TF expression, scRNA-seq analysis revealed that the majority of hypothalamic subtypes were generated in the *Eed-cKO* mutants. However, there was an increase in Glutamatergic/GABAergic double-positive cells, as well as loss/reduction of dopamine, Hypocretin/Orexin and a sub-group of Tac2 cells. scRNA-seq analysis revealed that these effects may result from dysregulation of several known cell-fate determinants. These findings suggest that many, but not all, aspects of the gene regulatory pathways necessary for hypothalamic development can play out irrespective of the H3K27me3 epigenetic mark, but points to higher sensitivity of certain neuronal sub-types to an altered epigenomic landscape.

## RESULTS

### Conditional knock out of *Eed* in the CNS results in the loss of H3K27me3

Constitutive *Eed* mutants die during early embryogenesis (Faust et al., 1998; Faust et al., 1995; Schumacher et al., 1996). To circumvent this early lethality we knocked out *Eed* conditionally in the CNS, by crossing a previously generated floxed *Eed* allele (Xie et al., 2014) to *Sox1-Cre* (Takashima et al., 2007). *Sox1* is expressed in the entire CNS and commences in the neural plate at E7.5 (Pevny et al., 1998). *Sox1-Cre* is a *Cre* insertion into the *Sox1* locus, and expresses *Cre* in agreement with the *Sox1* gene, as evident by Cre-mediated activation of a *ROSA26R-EYFP* reporter strain already at E8.5 in the entire developing CNS (Takashima et al., 2007). In contrast to *Eed* constitutive mutants, our conditional *Eed* mutants (denoted *Eed-cKO*) developed until at least E18.5. Hence, deleting *Eed* using *Sox1-Cre* circumvents early lethality while specifically removing *Eed* function from the entire developing hypothalamus at the earliest possible stage.

We previously found that *Eed-cKO* embryos displayed loss of H3K27me3 immunostaining in the dorsal telencephalon and lumbo-sacral spinal cord, with a reduction of immunostaining at E10.5 and a complete loss at embryonic day E11.5 (Yaghmaeian Salmani et al., 2018). Focusing upon the developing hypothalamus we observed a loss of H3K27me3 immunostaining in the hypothalamus in *Eed-cKO* mutants, at E11.5 (Figure 1SA-J). Based upon these findings, and our previous study, we conclude that deletion of *Eed* by the early CNS-specific deleter *Sox1-Cre* results in loss of the H3K27me3 mark, albeit with a 2-3 day delay.

### *Eed-cKO* mutants display reduced hypothalamic proliferation

*Eed-cKO* mutants display reduced proliferation in the dorsal telencephalon, while the lumbo-sacral spinal cord did not display any apparent change in proliferation (Yaghmaeian Salmani et al., 2018). To address proliferation in the hypothalamus, we used the CNS progenitor marker Sox2, phosphorylated Ser-28 on Histone 3 (PH3, a marker of mitotic cells), and 4’,6-diamidino-2-phenylindole (DAPI) nuclear staining, to assess total nuclear (cellular) volume. Focusing on E15.5, in the *Eed-cKO* mutants we observed fewer PH3+ cells in the hypothalamus than in control (*Eed^fl/fl^*) (Figure S2A-H). Quantification supported this notion and revealed significantly fewer PH3+ cells/DAPI volume and fewer PH3+ cells/Sox2 volume (Figure S2I-J). We did not however observe reduction of the percentage of Sox2 expressing cells/DAPI volume (Figure S2K). Hence, similar to our previous findings for the telencephalon (Yaghmaeian Salmani et al., 2018), *Eed-cKO* mutants displayed reduced proliferation in the hypothalamus during embryogenesis.

### Single cell transcriptional profiling reveals upregulation of H3K36me3 marked genes in *Eed* mutants

To address the effects of *Eed* mutation upon hypothalamus development in more detail we performed scRNA-seq analysis. We focused first upon a late embryonic stage, E18.5, because the majority of distinct neuronal sub-types – such as those specifically expressing neuropeptides and neurotransmitters – have been generated by this stage (Kim et al., 2020; Romanov et al., 2020; Zhang et al., 2021). We dissected the hypothalamus from 6 control (*Eed^fl/fl^*) and 9 *Eed-cKO* embryos, at E18.5. Cells were dissociated and sequenced on the Illumina/Bio-Rad ddSEQ platform, yielding a total of 20,703 cells.

Based upon the distribution profile of unique molecular identifiers (UMIs) and UMIs/gene, we removed cells with fewer than 600 UMIs and less than 1.2 UMIs/expressed gene. The remaining 14,121 cells had a mean of 5,275 UMI counts/cell and 1.9 counts/expressed gene (gFigure S3). Based on recent scRNA-seq analysis of the entire adult mouse nervous system (Zeisel et al., 2018) we identified non-neural cells e.g., vascular cells, blood cells and microglia, and because these would not have been targeted by *Sox1-Cre* (Yaghmaeian Salmani et al., 2018), we excluded them from further analysis. Moreover, *Foxg1* is expressed in the telencephalon and since its expression abuts the anterior extent of the hypothalamus (Figure 1A), these cells were also excluded. This yielded 11,713 hypothalamic neural cells, denoted “All-Hypo”; 6,202 cells from control and 5,511 cells from *Eed-cKO* embryos (Figure 1B). Uniform Manifold Approximation and Projection (UMAP) embedding, based upon the top-300 most variable genes (see Experimental Procedures; Table S1), illustrate this filtering process (Figure 1C).

**Figure 1.**
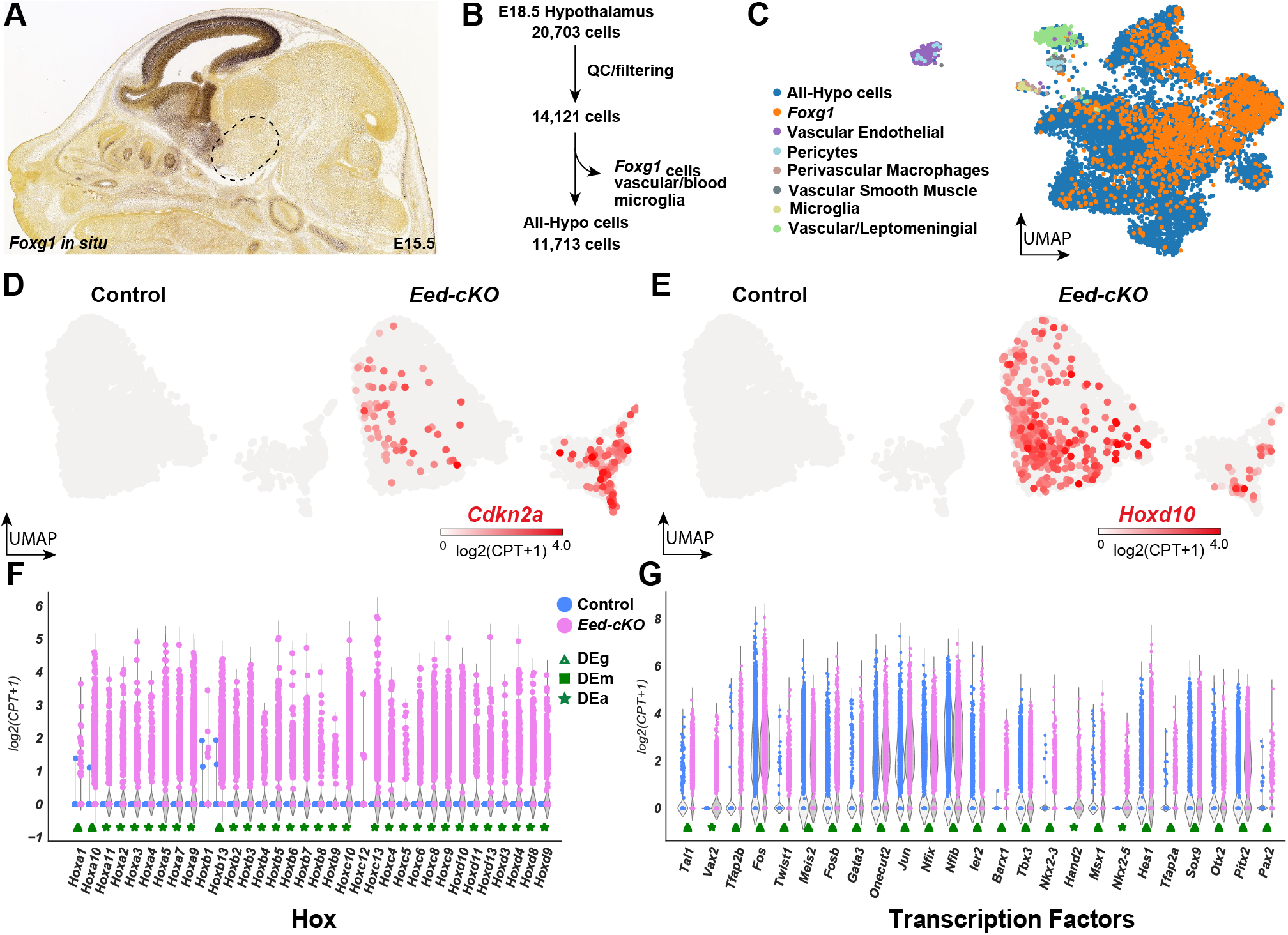
Single cell transcriptomic analysis of *Eed-cKO* mutants reveals upregulation of TFs and Hox genes in the hypothalamus. (A) *In situ* hybridisation of *Foxg1* in cross-section of E15.5 mouse embryo (image from Allen Brain Atlas) (Thompson et al., 2014). The dissected region analysed by scRNA-seq is boxed. (B) Process for quality control and biological filtering of hypothalamic scRNA-seq cells. (C) UMAP of hypothalamic scRNA-seq cells, combined from control and *Eed-cKO*, at E18.5, based upon 300 genes (Table S1). (D-E) UMAP of E18.5 hypothalamic scRNA-seq All-Hypo cells, showing expression of *Cdkn2a* and *Hoxd10*, in control and *Eed-cKO*. (F-G) Violin plots comparing the expression levels of Hox genes (F) and the highest differentially expressed (adjusted p-value ≤ 0.05) transcription factors (G) between control and *Eed-cKO*, in E18.5 hypothalamic scRNA-seq All-Hypo cells. Markers below violin plots denote differentially expressed genes; green markers indicate up-regulation in *Eed-cKO*. The shape of markers demarcates the type of differential expression; triangles indicate genes which are generally differentially expressed (DEg, differential expression), squares indicate genes which change in the magnitude of expression (DEm, differential magnitude), and stars indicate genes which change in the number of cells expressing the gene (DEa, differential abundance).

To assess differential gene expression (DE) we used *DEsingle* (Miao et al., 2018). We categorized the effects as: differential abundance of cells expressing the gene (DEa), differential magnitude of gene expression (DEm), or general differential expression (DEg), which entails both differential abundance and magnitude. Analysing the All-Hypo cells revealed 5,421 DE genes, when comparing *Eed-cKO* to control (Table S2); 2,307 genes were upregulated and 3,114 were down-regulated. Of the upregulated genes, we found that 104 genes were DEa, 1 DEm, and 2,202 were DEg; while for the downregulated genes 11 were DEa, 0 DEm, and 3,103 DEg. Analysis for annotated gene sets enriched in our groupings of DE genes revealed an enrichment for genes involved in transcriptional regulation and H3K27me3 marked genes across the *ENCyclopaedia of DNA Elements* (*ENCODE*) (Davis et al., 2018). In addition, the upregulated DEg group also showed enrichment for H3K4me3 and H3K36me3 marked genes (Figure S4, Tables S3 and S4).

### Single cell transcriptional profiling reveals cell cycle inhibitor upregulation and brain posteriorization in *Eed* mutants

Our previous study of *Eed-cKO* mutants revealed reduced proliferation in the developing telencephalon, accompanied by upregulation of two members of the Cip/Kip family of cell cycle inhibitors; *Cdkn1a* and *Cdkn1c* (Yaghmaeian Salmani et al., 2018). In line with these findings, analysing the All-Hypo cells, using UMAP embedding and DE analysis, revealed upregulation of *Cdkn1b* and *Cdkn1c* (both DEg), as well as of three of the four members of the INK4 family of cell cycle inhibitors, *Cdkn2a* (DEa), *Cdkn2b* (DEg) and *Cdkn2c* (DEg) (Figure 1D, Table S2).

Turning to developmental regulatory genes, UMAP embedding and DE analysis revealed that 33 out of the 39 Hox homeotic genes were upregulated in the *Eed-cKO* mutants (3 DEg, 29 DEa) (Figure 1E-F, Table S2). This is in line with our previous analysis of the telencephalon, using bulk RNA-seq, which showed upregulation of all 39 Hox genes, to levels matching their expression in the spinal cord (Yaghmaeian Salmani et al., 2018). Moreover, we observed that many other posteriorly expressed TFs were also upregulated, including *Pax2*, which displayed upregulation in the UMAP embedding, in the DE analysis (DEg) and on immunostaining (Figure 1G, S5A-F, Table S2). Hence, the scRNA-seq and immunostaining analysis revealed that *Eed-cKO* mutants displayed posteriorization and a reduced proliferative gene expression profile in the developing hypothalamus.

### Single cell transcriptional profiling identified all major hypothalamic cell types at E18.5

The ectopic expression of a number of posteriorly expressed TFs suggested that hypothalamic cell specification may be strongly affected in the *Eed-cKO* mutants. To address this issue, we first analysed the generation of broader hypothalamic cell types (Chen et al., 2017; Kim et al., 2020; Romanov et al., 2020; Romanov et al., 2017). UMAP embedding of All-Hypo cells revealed that the major cell types were generated in *Eed-cKO* mutants, including GABAergic neurons (*Slc32a1*), glutamatergic neurons (*Slc17a6*), oligodendrocytes (*Olig1*), astrocytes (*Gfap*), ependymal cell (*Foxj1*) or tanycytes (*Rax*) (Figure 2A-F). Nevertheless, DE analysis did reveal minor downregulation of *Slc32a1* (DEg) and *Slc17a6* (DEg), while *Olig1* (DEg), *Gfap* (DEg), *Rax* (DEa) and *Foxj1* (DEg) were upregulated (Table S2). Moreover, we observed an apparent increase in the overlap between glutamatergic and GABAergic markers (*Slc32a1* and *Slc17a6*) (Figure 2E-F). Specific identification of double-positive cells confirmed this observation, revealing a significant increase in double-positive cells (Figure 2G, Fisher’s Exact Test p-value < 1e-3). This effect prompted us to scrutinize the TF expression changes in the *Eed-cKO*. We noted that the *Tfap2a* and *Tfap2b* genes, known determinants of cerebellar GABAergic interneuron cell fate (Zainolabidin et al., 2017), were among the genes ectopically expressed (Figure 1G). This was apparent both in the UMAP embedding and the DE analysis (both DEg) (Figure 1G, 3E-F, Table S2). Immunostaining confirmed this upregulation. Specifically, in control, Tfap2b is restricted to posterior regions of the E18.5 brain. In contrast, in *Eed-cKO*, Tfap2b expression was apparent throughout the entire brain, including the hypothalamus (Figure 3A-D).

**Figure 2.**
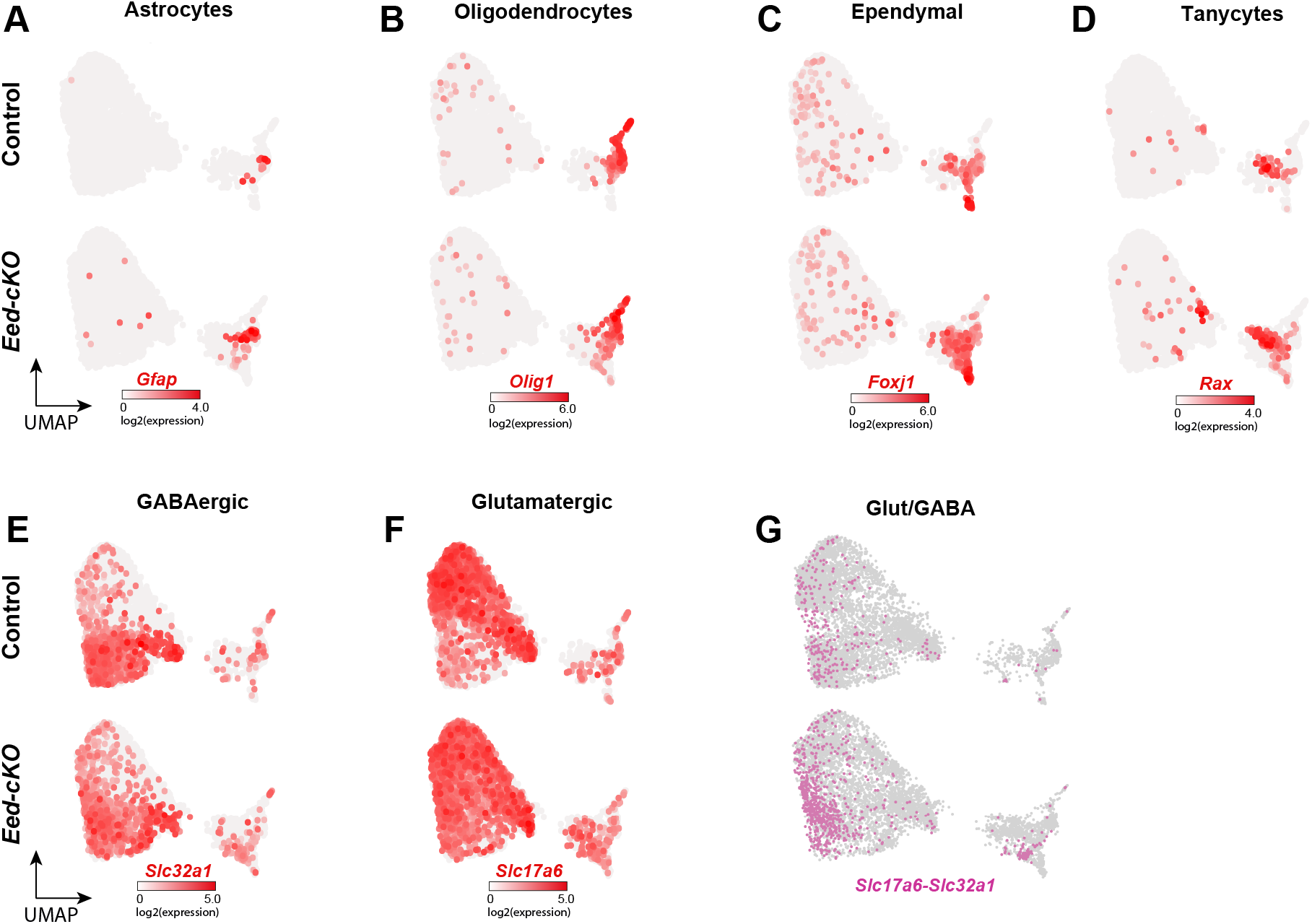
*Eed-cKO* mutants display more Glut/GABA cells. (A-F) UMAP embedding, based upon 300 DE genes (Table S1), of E18.5 All-Hypo cells, showing astrocytes (*Gfap*), oligodendrocyte precursors (*Olig1*), ependymal cells (*Foxj1*), tanycytes (*Rax*), GABAergic neurons (*Slc32a1*) and glutamatergic neurons (*Slc17a6*). (G) Glut/GABA co-expression revealed by *Slc32a1* and *Slc17a6* expression.

**Figure 3.**
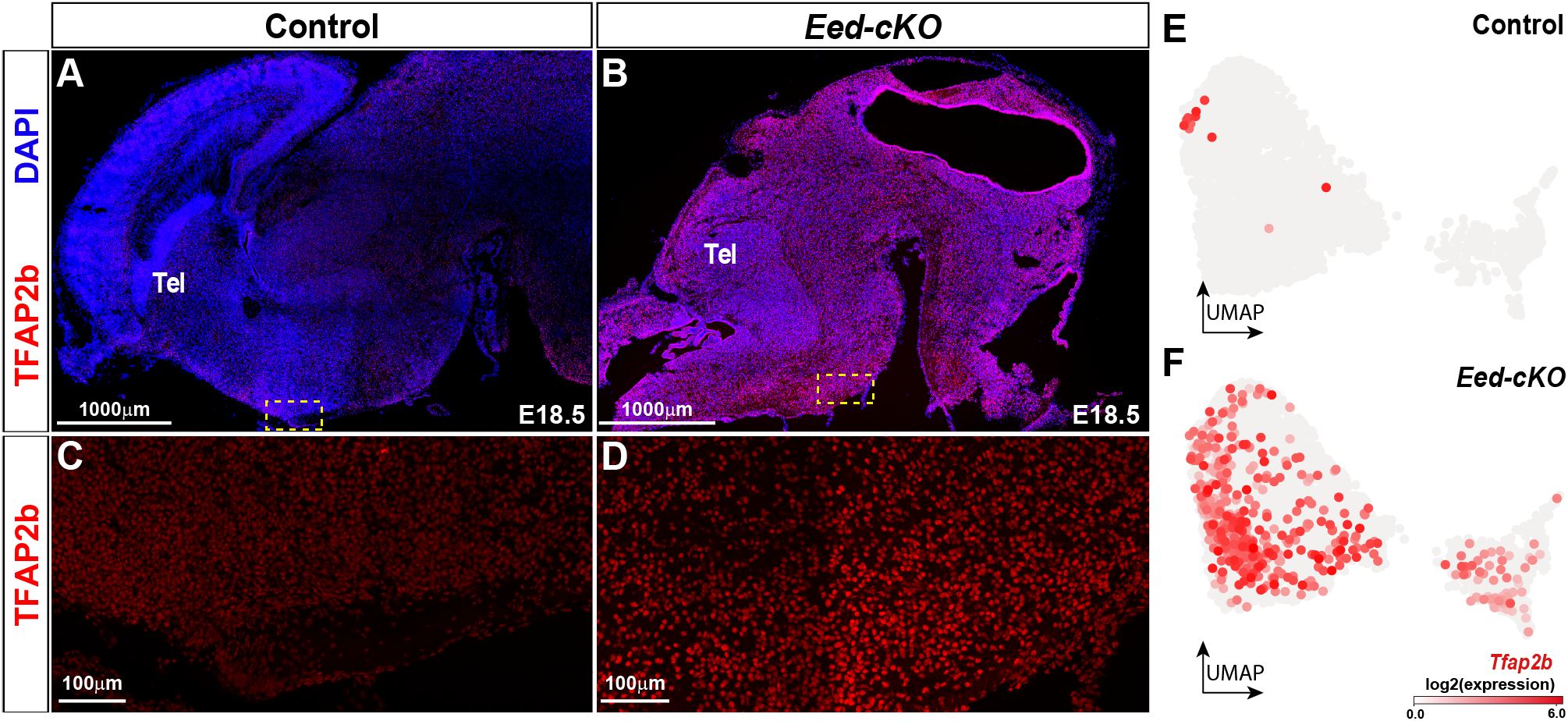
*Eed-cKO* mutants display ectopic Tfap2b expression in the hypothalamus. (A-D) Staining for DAPI and Tfap2b in sagittal sections of E18.5 brains, in control and *Eed-cKO*. Dashed squares in (A-B) delineate regions of hypothalamic tissue magnified in (C-D). In control, Tfap2b expression is observed in the mid-and hindbrain regions. In *Eed-cKO* mutants, Tfap2b expression is increased and expands into the anterior brain, including the hypothalamus. (E-F) UMAP embedding of E18.5 hypothalamic scRNA-seq All-Hypo cells, with each cell coloured according to the expression level of *Tfap2b* in control and *Eed-cKO*. Scale bar; 1,000 µm in (A, B), 100 µm in (C, D).

Hence, while all major hypothalamic sub-types were generated in *Eed-cKO*, there were effects on the partition of GABAergic and glutamatergic cell fate, and ectopic expression of the *Tfap2a* and *Tfap2b* GABAergic determinants.

### Single cell transcriptional profiling reveals limited effects on hypothalamic cell fate specification in *Eed* mutants

Next, we focused on the generation of more discrete neuronal subtypes. We analysed the All-Hypo cells for their expression of 29 neuropeptide (NP) genes (Table S1), the *Th* and *Ddc* dopamine markers (DA), and the *Hdc* histaminergic marker, and thereby identified 8,219 “NP-DA” cells within the All-hypo cells; 4,275 from control and 3,944 from *Eed-cKO* embryos. We created UMAP embedding of the NP-DA cells using the aforementioned NP-DA genes, as well as 90 additional genes found to be most correlated with the expression of the NP-DA marker genes (Table S1). In control, clustering and UMAP visualisation based upon these marker genes identified 116 major cell clusters (Figure 4A, 4C). Several clusters displayed co-expression of neuropeptides i.e., *Agrp-Npy*, *Avp-Gal*, *Kiss1-Tac2* and *Npy-Sst*, which is in agreement with previous scRNA-seq expression analysis in the adult and/or embryonic hypothalamus (Chen et al., 2017; Romanov et al., 2020; Romanov et al., 2017; Zhang et al., 2021). Expression of *Th* and *Ddc* is broader than DA neurons (Bjorklund and Dunnett, 2007), and *Th*-*Ddc* co-expression is only detected in two smaller cell clusters, which also express *Slc6a3*, identifying them as *bona fide* DA neurons (Figure 4A, 4C). Two neuropeptide genes (*Galp*, *Prlh*) were not detected.

**Figure 4.**
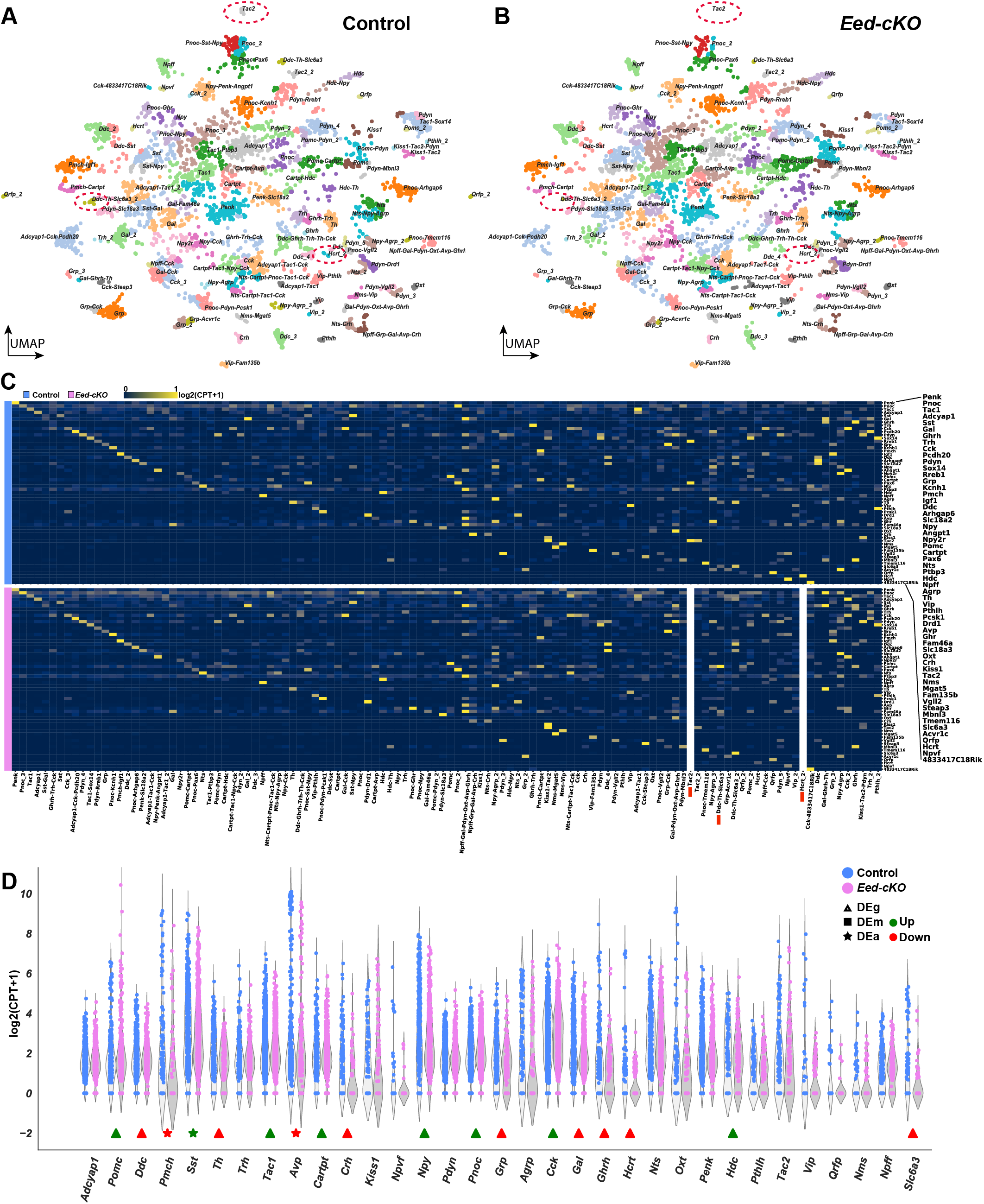
*Eed-cKO* is necessary for DA, Hcrt and Tac2 cells. (A-B) UMAP embedding of hypothalamic NP-DA cells, based on 122 genes (Table S1), in control and *Eed-cKO*, at E18.5. Each cluster is labelled by the combination of NP-DA marker genes upregulated in that cluster. (C) Heatmap displaying average gene expression of the NP-DA marker genes in each UMAP cell cluster, in control (blue) and *Eed-cKO* (pink). Gene expression is measured in log2(CPT+1), scaled between 0 and 1 along each row (see Mat&Met). The *Ddc-Th-Slc6a3*, *Tac2* and *Hcrt* clusters are absent in *Eed-cKO*, and hence a column of 0 expression was depicted. (D) Expression levels of NP-DA marker genes, in control and *Eed-cKO*, at E18.5. Symbols below violin plots indicate differentially expressed genes; green markers show up-regulation in *Eed-cKO* and red indicates down-regulation. The shape of markers demarcates the type of differential expression; triangles indicate genes which are generally differentially expressed (DEg, differential expression, general), squares indicate genes which change in the magnitude of expression (DEm, differential magnitude), and stars indicate genes which change in the number of cells expressing the gene (DEa, differential abundance). See Table S2 for analysis.

Turning to the *Eed-cKO* mutants, in spite of the reduced proliferation and the ectopic expression of a number of posterior TFs, the UMAP embedding revealed surprisingly limited effects upon NP-DA cell specification in the E18.5 hypothalamus, as evidenced by the generation of the majority of the 116 NP-DA sub-types (Figure 4B-C). Specification of NP-DA sub-types occurred in spite of the ectopic expression of Hox and other regulatory genes within the majority of cell clusters (Figure S6A-B). For example, in control, *Avp* expressing cells separated into four clusters, all of which co-expressed other neuropeptide genes. All four Avp cell clusters were present in the UMAP from *Eed-cKO* mutants, in spite of the ectopic expression of Hox genes and other TFs in these cells (Figure 4A-D, S6A-B, Table S2). The generation of Avp neurons in *Eed-cKO* was also supported by immunostaining (Figure S7A-H).

### Single cell transcriptional profiling reveals loss of DA neurons in *Eed* mutants

While the scRNA-seq and immunostaining analysis revealed that most NP-DA cell sub-types were generated in *Eed-cKO*, we noted several specific effects. Interestingly, UMAP embedding revealed that one of the *Ddc-Th-Slc6a3* clusters was largely lost in *Eed-cKO* mutants, while the other was reduced in size (Figure 4A-B). In line with this finding, *Ddc*, *Th* and *Slc6a3* were all significantly downregulated in *Eed-cKO* mutants (DEg) (Figure 4D, Table S2, Figure 5A-C). Studies of hypothalamic DA cells have revealed the involvement of several TFs, including Rax, Dlx1, Otp, Sim1 and Arnt2 (Biran et al., 2015; Orquera et al., 2016; Yee et al., 2009). Because the *Ddc-Th-Slc6a3* cells are largely missing in *Eed-cKO*, we could not determine the expression of these genes in the DA cells. However, DE analysis of the All-Hypo cells revealed that *Dlx1* and *Arnt2* were both downregulated in All-Hypo cells (both DEg), while, surprisingly, *Rax* was upregulated in All-Hypo cells (DEg) (Table S2). *Otp* was neither detected in control nor *Eed-cKO* (Table S2).

**Figure 5.**
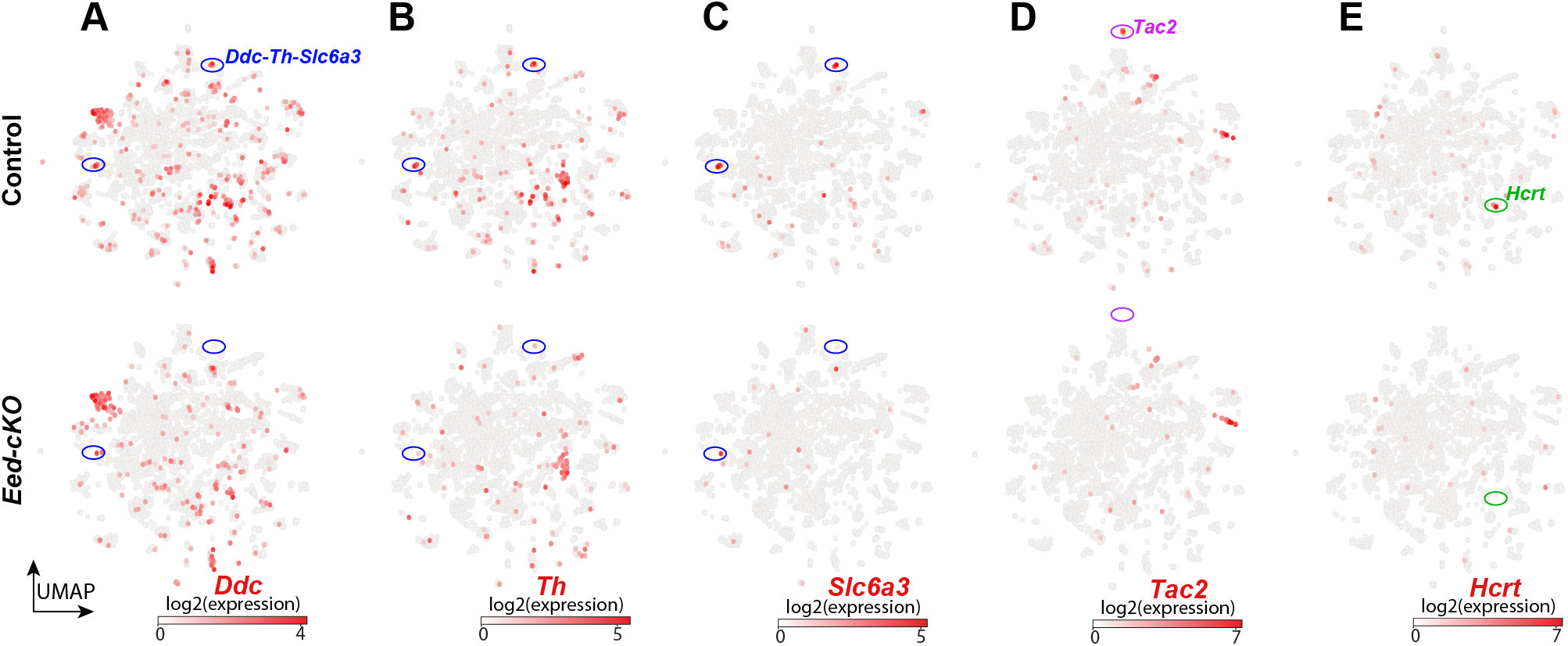
*Eed-cKO* is necessary for DA, *Hcrt* and *Tac2* cells. (A-E) UMAP embedding of E18.5 NP-DA cells, based upon 122 DE genes (Table S1), showing expression of *Ddc*, *Th*, *Slc6a3*, *Tac2* and *Hcrt*. (A-C) Two prominent clusters co-express *Ddc*-*Th*-*Slc6a3*, and these cluster are largely missing in *Eed-cKO*. (D-E) Similarly, one specific *Tac2* cluster and the *Hcrt* cluster are largely absent in *Eed-cKO* mutants.

### Single cell transcriptional profiling reveals loss of Hcrt neurons in *Eed* mutants

Turning to neuropeptide cells, while UMAP embedding did not reveal obvious effects in *Eed-cKO* upon the majority of neuropeptide genes, DE analysis did reveal minor effects upon a number of them, including upregulation of 8 genes and downregulation of 7 genes (Figures 4D, 5D-E, S8, S9, Table S2). In contrast, for *Tac2*, while UMAP analysis revealed a complete loss of one *Tac2*-only cluster, this was not confirmed in the DE analysis, probably due to *Tac2* being expressed in three other clusters, which appeared unaffected in the UMAP (Figures 4A-B, 4D, 5D). The upstream regulatory cues specifying the different Tac2 neuronal sub-types is not well understood, precluding any assessment of potential causes of the loss of one specific Tac2 cluster.

We also observed strong effects upon *Hcrt*, both with respect to a near complete loss of cells expressing *Hcrt* in the UMAP, and to the expression levels (DEg) (Figure 4A-D, 5E, 6G-H, Table S2). This observation was confirmed by immunostaining (Figure 6A-D). Previous studies have identified several TFs that are involved in Hcrt cell specification, including Dbx1, Ebf2 and Lhx9 (Dalal et al., 2013; De La Herran-Arita et al., 2011; Liu et al., 2015; Sokolowski et al., 2015). In control *Hcrt* cells, *Lhx9* is the only TF gene of these three that is DE in *Hcrt* cells, when compared to other NP-DA clusters (Table S5). Because the *Hcrt* cells are missing in *Eed-cKO*, we could not determine the expression of *Dbx1*, *Ebf2* or *Lhx9* in these cells in the mutant. However, DE analysis of All-Hypo cells revealed that both *Ebf2* and *Lhx9* were downregulated (both DEg) (Table S2).

**Figure 6.**
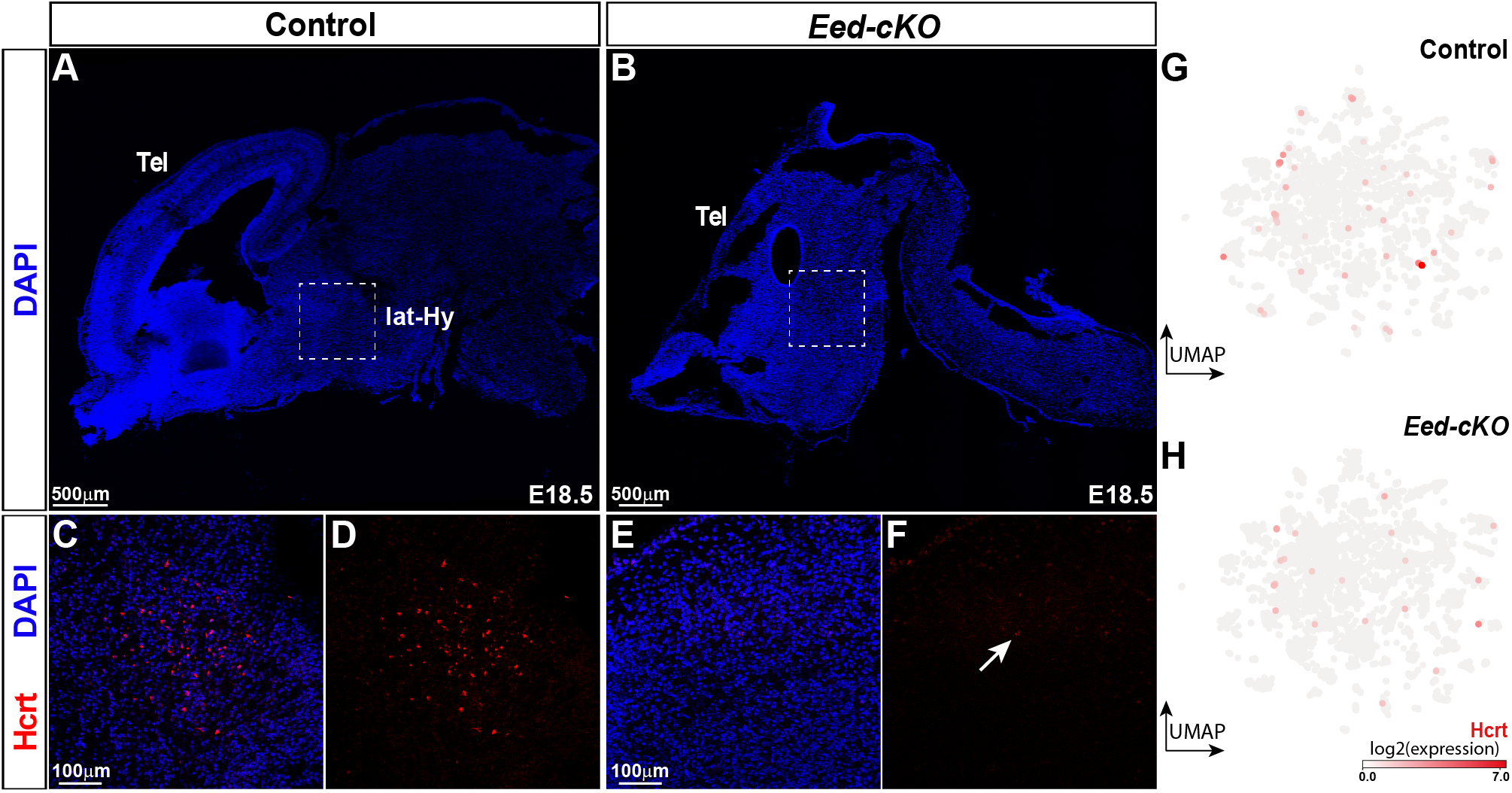
*Eed* is critical for Hcrt neurons. (A-B) Staining for DAPI in sagittal sections of E18.5 control and *Eed-cKO* mutant brains. Dashed, square insets delineate the lateral hypothalamus. (C-F) Staining for DAPI and Hcrt in the lateral hypothalamus, in control and *Eed-cKO* embryos. A cluster of Hcrt cells is present in control, while *Eed-cKO* mutants only show occasional, weakly expressing cells. (G-H) UMAP embedding of E18.5 hypothalamic scRNA-seq NP-DA cells, with each cell coloured according to the expression level of *Hcrt* in control and *Eed-cKO*. Scale bar; 500 µm in (A, B), 100 µm in (C-F).

### scRNA-seq at earlier stages reveals reduction of Lhx9 expressing progenitor cells in *Eed* mutants

To better understand the origins of the phenotypes observed at E18.5 we conducted scRNA-seq analysis of control and *Eed-cKO* at earlier stages; E13.5, E15.5 and E18.5. We generated a total of 51,540 high-quality cells: 21,335 from control and 30,205 from *Eed-cKO* mutants. Following the same biological filtering process as for E18.5, we yielded a total of 41,646 hypothalamic neural cells (All-Hypo); 18,551 cells from control and 23,095 cells from *Eed-cKO* embryos (Figure S10A-C). Additional stages from control were also generated to capture the transcriptional range of phenotypes across hypothalamic development for determining the UMAP embedding, though only equivalent stages were analysed for all subsequent statistical analysis (see Methods).

First, we focused first on the three NP-DA cell types affected at E18.5: the DA (*Slc6a3*), *Hcrt* and *Tac2* sub-cluster of cells. In control, all three cell types were evident also at the earlier embryonic stages. However, in *Eed-cKO* mutants there was a near complete loss of *Slc6a3* expressing cells, a complete loss of high-expressing Hcrt cells, and loss of one *Tac2* cluster across all stages (Figures 7E, S10C). Hence, all three NP-DA cell types that were affected at E18.5 were also affected at earlier stages.

**Figure 7.**
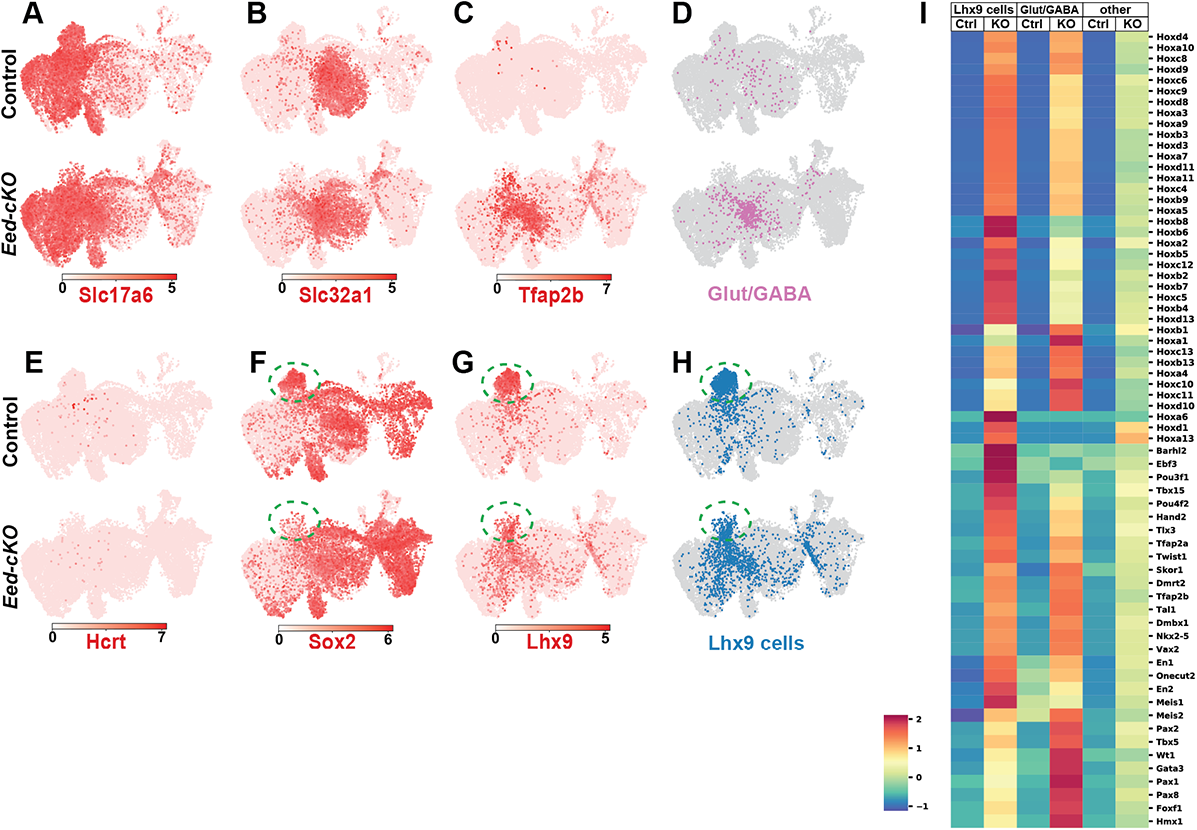
Developmental onset of Glut/GABA co-expression and loss of Lhx9/Sox2 progenitors. (A-H) UMAP embedding of hypothalamic cells from E13.5, E15.5 and E18.5. (A-D) Glut/GABA co-expression and Tfap2 expression is limited in control but increases in *Eed-cKO* mutants. (E) High-expression Hcrt cells are not present in *Eed-cKO* mutants. (F-H) In control, a set of *Sox2/Lhx9* progenitor cells are present (green dashed oval), but this cell group is greatly reduced in *Eed-cKO* mutants. (I) Expression analysis of *Lhx9* cells, Glut/GABA cells and “other” cells, at E13.5-E18.5. In *Eed-cKO*, *Lhx9* and Glut/GABA cells show ectopic expression of a number of developmental TFs, and display upregulation to a greater extent than all “other” cells.

To address broader aspects of development we utilised a binary labelling method with cell type marker genes (see Methods), according to broad cell types such as Glutamatergic, GABAergic, sub-ventricular zone, ventricular zone, and various glial cell types (Figure S10C). We noticed two major differences between control and *Eed-cKO*. First, the loss/reduction of a particular subpopulation of progenitor cells (Hes5-, Sox2+) (Figures 7F, S10C). Because Lhx9 is one of the few known transcription factors involved in *Hcrt* cell specification and was significantly downregulated in *Eed-cKO* at E18.5, we examined *Lhx9* expression in the temporal UMAP space. This revealed that the specific Hes5-, Sox2+ progenitor population that displayed loss/reduction in *Eed-cKO* also co-expressed *Lhx9* (Figure 7F-H). Gene expression analysis of the *Lhx9* group of cells between control and *Eed-cKO*, as well as against all mutant cells (Table S9, S10) revealed significant upregulation of many developmental regulators in the *Lhx9* population, including 29/39 of the Hox transcription factors (Figure 7I, Table S11). Second, similar to what was observed at E18.5 we noted significantly higher numbers of Glut/GABA cells in *Eed-cKO* mutants (Figure 7A-D, Fishers exact test, p<1e-3). Again, gene expression analysis revealed ectopic expression of developmentally important genes in the Glut/GABA cells, such as *Tfap2a/b* and *Pax2* (Figure 7C,I, Table S11). We observed significantly greater ectopic expression of developmental regulators in both the *Lhx9* and Glut/GABA cells, when compared to all other cells (Figure 7I, Table S11), revealing that these two cell populations were particularly affected.

In summary, *Eed-cKO* mutants display a loss of the H3K27me3 mark from E11.5 and onward, posteriorization of the hypothalamus, and reduced proliferation. Despite these effects most major neuronal sub-types are generated by E18.5. However, there is an increase in the number of Glut/GABA co-expressing cells, and loss/reduction of DA, *Hcrt* and a sub-set of *Tac2* neurons. These effects are likely due to the up-and downregulation of several known upstream regulators in specific subpopulations of progenitors. Interestingly, despite the gradual loss of H3K27me3, we did not observe any apparent correlation between the specific cells affected and the onset of their marker genes (Table S6).

## DISCUSSION

### Gradual loss of H3K27me3 in *Eed* mutants

Epigenomic methylation marks are generally viewed as stable over time. However, they are subjected to two main removal cues: de-methylation and replication-mediated dilution. How do these removal dynamics fit with the observed gradual reduction of H3K27me3 in *Eed-cKO*?

De-methylation of H3K27me3 is chiefly mediated by *Kdm6a* (*Utx*) and *Kdm6b* (*JMJD3*) (Arcipowski et al., 2016). However, while mutation of *Kdm6a* and/or *Kdm6b*, in mouse or ESCs, did result in elevated H3K27me3 levels at specific loci, there was no global increase in H3K27me3 levels (Burgold et al., 2008; Park et al., 2014; Tang et al., 2020; Wijayatunge et al., 2018). Similarly, in *Drosophila*, *Utx* mutation did not alter the dynamics of H3K27me3 reduction at a specific target locus in the developing wing disc (Laprell et al., 2017). Hence, while de-methylation mediated by Kdm6a/b modulates H3K27me3 levels, it does not appear to play an instrumental role in counteracting PRC2 activity at all target loci.

By contrast, DNA replication can efficiently reduce the levels of the H3K27me3 mark by H3 dilution, as new nucleosomes form post replication. Studies in *Drosophila* have found that removal of a Polycomb Response Element (PRE) at a target locus resulted in the gradual reduction of H3K27me3, with some 50% reduction per cell cycle, and loss of transcriptional repression already after one cell cycle (Coleman and Struhl, 2017; Laprell et al., 2017). Similarly, studies of the mouse intestine have revealed that loss of PRC2 activity (by *Eed* or *Ezh2* mutation) resulted in the gradual loss of H3K27me3 levels, also with an estimated 50% reduction per cell cycle (Jadhav et al., 2020).

To circumvent early embryonic lethality we inactivated *Eed* using the early neuroectodermal Cre deleter, *Sox1-Cre*, (active at E8.5) (Takashima et al., 2007). This resulted in a gradual loss of the H3K27me3 mark in the *Eed-cKO* mutants, with reduction in immunostaining at E10.5 and complete loss at E11.5, in the telencephalon, spinal cord and hypothalamus (Yaghmaeian Salmani et al., 2018) (herein). We envision that some of the delay in H3K27me3 reduction results from the delay in deletion of both gene copies, as well as the degradation of wild type *Eed* mRNA and Eed protein, the latter of which may be especially slow for Eed protein that is associated with the PRC2 complex. In addition, a major part of the gradual reduction of H3K27me3 levels is likely connected to proliferation dynamics. Progenitor cycling speeds have not been mapped in the early mouse hypothalamus, but studies of the E11 mouse telencephalon has revealed neuroepithelial progenitor cell cycle speeds of 8-10 hrs (Caviness et al., 1995; Takahashi et al., 1995). In light of these studies, the gradual loss of the H3K27me3 mark in *Eed-cKO* mutants, being reduced at E10.5 and lost at E11.5 i.e., 2-3 days and hence some 6-9 cell divisions after *Eed* inactivation at E8.5, is consistent with a replication-mediated dilution of modified H3 histones.

### Reduced proliferation in *Eed-cKO* mutants

Previous studies of *Eed-cKO* mutants revealed reduced proliferation in the telencephalon (Telley et al., 2019; Yaghmaeian Salmani et al., 2018), which was accompanied by increased expression of the Cip/Kip cell cycle inhibitors *Cdkn1a* and *Cdkn1c* (Yaghmaeian Salmani et al., 2018). Here, we observed reduced proliferation also in the hypothalamus, and upregulation of *Cdkn1b* and *Cdkn1c*, as well as *Cdkn2a*, *Cdkn2b*, and *Cdkn2c*, in the E18.5 scRNA-seq data. It is likely that the upregulation of most of the Cip/Kip and INK4 cell cycle inhibitor genes is a major reason for the reduced proliferation in *Eed-cKO* mutants.

The reduced proliferation in the hypothalamus begs the question of which cells are not being generated. Recent scRNA-seq analysis of the developing hypothalamus has revealed that neurons are generally born earlier than other cell types, appearing already at E10 and onward, and that glia, ependymal cells and tanycytes appear from E16 and onward (Kim et al., 2020; Romanov et al., 2020; Zhang et al., 2021). Our UMAP embedding of the All-Hypo cells did not reveal striking loss of any major cell type i.e., GABAergic neurons, glutamatergic neurons, astrocytes, oligodendrocytes, ependymal cells and tanycytes. However, we observed an increase in Glu/GABA co-expressing cells. Single expressing Glu and GABA cells, as well as Glu/GABA co-expressing cells, are born early during hypothalamic development, but are also added throughout embryogenesis [herein (Kim et al., 2020; Romanov et al., 2020; Zhang et al., 2021)]. Against this backdrop, it is not apparent if/how the reduced proliferation observed in *Eed-cKO* mutants could act to increase the number of Glu/GABA co-expressing cells.

Regarding NP-DA subtypes, while the actual birth-date for most hypothalamic neuronal subtypes has not been determined, the onset of NP-DA marker expression has been analysed in most cases (Table S6). Assuming a gradual loss of the H3K27me3 mark, a logical effect would be the loss of late-expressing NP-DA markers, indicating a failure to generate late-born neurons. However, the subtypes affected do not apparently link to marker gene expression onset, with genes activated both early (E11.5) and late (E15.5) being affected, or alternatively, unaffected (Table S6). Hence, the differential effects upon discrete cell subtypes are not readily explained by differential birth dates.

Previous analysis in the telencephalon revealed that *Eed-cKO* mutation primarily affected the proliferation of daughter cells, as opposed to progenitor cells (Yaghmaeian Salmani et al., 2018). Recent studies, including genetic lineage tracing, have revealed a complex repertoire of progenitor and proliferative daughter cells in both the mouse and human hypothalamus, which in many aspects are similar to the developing telencephalon (Zhang et al., 2021; Zhou et al., 2020). It is tempting to speculate that the selective effects of *Eed-cKO* upon cellular subtypes born both early and late may relate to the lineage topology of different hypothalamic lineages, an issue of which we still know very little.

### *Eed-cKO* triggers posteriorization of the brain

A well-conserved role of PRC2 in Bilaterians is to confine Hox homeotic gene expression to the posterior CNS (Isono et al., 2005; Li et al., 2011; Struhl, 1983; Struhl and Akam, 1985; Suzuki et al., 2002; Wang et al., 2002; Yaghmaeian Salmani et al., 2018). Our previous studies of *Eed-cKO* mutants confirmed this role for PRC2, and we observed ectopic expression of all 39 Hox homeotic genes in the E15.5 mouse forebrain, as revealed by bulk RNA-seq (Yaghmaeian Salmani et al., 2018). We also observed several other, posteriorly expressed TFs being ectopically expressed in the forebrain. Here, using scRNA-seq, we find similar effects in the developing hypothalamus. A growing body of work is pointing to the role of Hox genes in regulating cell cycle genes and repressing proliferation (Economides et al., 2003; Monedero Cobeta et al., 2017; Yaghmaeian Salmani et al., 2018). Strikingly, in *Drosophila*, simultaneous removal of Hox genes can rescue reduced proliferation in *esc* mutants (the *Drosophila* orthologue of *Eed*) (Bahrampour et al., 2019). These studies indicate that the reduced hypothalamic proliferation observed in *Eed-cKO* may be a direct result of the posteriorization of the entire fore-and midbrain, and the ectopic expression of posterior Hox homeotic genes.

### Enrichment of H3K36me3 in genes repressed by PRC2

The recruitment of PRC2 to chromatin and the control of its local activity has been, and continues to be, under intense investigation. This issue is further complicated by the finding that different PRC2 sub-complexes have different properties. There are five major recruitment and/or activity interaction nodes for mammalian PRC2: low-methylated CpG islands, H2AK119ub, H3K27me3, H3K36me3 and RNA (Chammas et al., 2020; Laugesen et al., 2019; Mocavini and Di Croce, 2020; van Mierlo et al., 2019). Recent studies found that activation of genes repressed by PRC2, post *Eed* or *Ezh2* mutation, was positively correlated with the levels of not only H3K27me3, but also the H3K4me3 active mark (Jadhav et al., 2020). In line with these findings, our GO enrichment analysis also identified an enrichment of DE genes marked by H3K27me3 and H3K4me3. In addition, we observed enrichment of the H3K36me3 mark, specifically in the upregulated genes. While generally regarded as an activating mark, acting by repressing PRC2 docking and activity (Finogenova et al., 2020), H3K36me3 has also been proposed to selectively mark genes for future H3K27me3 modification and subsequent repression to facilitate transition from pluripotency to post-mitotic cell fates [reviewed in (Abed and Jones, 2012)]. Since the category “upregulated DEg genes” represent those that had both increased magnitude of expression and an increased number of cells expressing that gene, it is possible that this group of genes may represent those that should have been repressed by PRC2 to facilitate transition to post-mitotic cell fates.

### Selective involvement of *Eed* upon hypothalamic cell fates

In *Eed-cKO* mutants, despite the reduced proliferation and upregulation of many posteriorly expressed developmental regulators, we observed that the majority of neural sub-types were still generated. One possible explanation for the limited effects observed may be that *Eed-cKO* results in the loss of H3K27me3 subsequent to that many important developmental decisions, i.e., patterning of the hypothalamic progenitor region, have already played out, during E8-E11 (Alvarez-Bolado, 2019; Bedont et al., 2015; Blackshaw et al., 2010; Burbridge et al., 2016; Ferran et al., 2015; Puelles and Rubenstein, 2015; Xie and Dorsky, 2017). Nevertheless, major aspects of neurogenesis and cell specification do occur subsequent to E11.5, and while there is extensive mis-regulation of a large number of genes it is surprising that H3K27me3 is apparently not critical for many of the genetic cascades that govern cell fate.

Although many cell types are present, we did observe an increase in Glut/GABA co-expressing cells, apparent at all stages analysed. We observed DE of a number of TFs, including ectopic expression of the cerebellar GABA determinants *Tfap2a* and *-b* (Zainolabidin et al., 2017), and their expression showed extensive overlap with Glut/GABA co-expressing cells.

In addition, *Eed-cKO* mutants also displayed the absence of DA and Hcrt cells, as well as a subset of Tac2 cells. While the absence of these cells in *Eed-cKO* mutants hindered detailed analysis of the underpinnings of these phenotypes, we observed that two of the three known regulators of hypothalamic DA cells, *Dlx1* and *Arnt2*, were downregulated. Similarly, we observed down-regulation of two of the three known regulators of *Hcrt* cells, *Ebf2* and *Lhx9*. Down-regulation of these TF genes may indeed explain the absence of DA and Hcrt cells. Indeed, analysis of earlier stages revealed that a subset of *Lhx9* positive progenitors (Sox2+) are missing in the mutant. Specification of the different *Tac2* neuronal sub-types has not been previously addressed, and hence the underpinnings of this phenotype is currently unclear.

Are there any commonalities between DA, Hcrt and Tac2 neurons that could explain the absence of these three particular cell types? The specific progenitor region of origin for the different hypothalamic DA neuron sub-types (A11, A12, A14, A15), Hcrt or Tac2 neurons is not clear, but their final cell body locations (DA neurons medial, Hcrt neurons lateral and Tac2 neurons in several regions) argue against a common origin. Similarly, the non-overlapping sets of upstream regulators, Otp, Sim1 and Arnt2 for DA neurons (Biran et al., 2015), and Dbx1, Ebf2 and Lhx9 for Hcrt neurons (Dalal et al., 2013; De La Herran-Arita et al., 2011; Liu et al., 2015; Sokolowski et al., 2015), also argue against a common origin, as well as against a common genetic pathway. Interestingly, knock-out of the H3K4-methyltransferase *Mll4* gene in the developing hypothalamic arcuate nucleus also resulted in highly selective effects on neuronal sub-type specification (Huisman et al., 2021).

### Implications for PRC2 sensitivity

A growing body of work points to the role of epigenetics in gating entry into puberty, in particular with respect to controlling the expression of neuropeptide genes involved in triggering puberty, such as *Kiss1* and *Tac2* (aka *Tac3*) (Aylwin et al., 2019; Shalev and Melamed, 2020). This pertains specifically to the PRC2 complex, which plays a major role in gating the elevated expression of the *Kiss1* and *Tac2* genes necessary for puberty. Analysis of a number of PRC2 genes revealed that *Eed* gene expression is downregulated in the hypothalamus pre-puberty, followed by *Kiss1* and *Tac2* upregulation. Strikingly, overexpression of *Eed* resulted in repression of *Kiss1* gene expression and an inhibition/delay of puberty (Lomniczi et al., 2013). We did not, however, observe any upregulation of *Kiss1* or *Tac2* in our *Eed-cKO* scRNA-seq data (Table S2), but rather the loss of a distinct cluster of *Tac2* expressing cells. These findings indicate that the epigenomic profile of the *Kiss1* and Tac2 promoters and their sensitivity to PRC2 may change from embryogenesis to early adult life, and from “baseline” to elevated levels of expression.

Finally, in humans, extensive genome and exome sequencing points to that all major PRC2 components, including *EED*, are haploinsufficient (Karczewski et al., 2020). In line with this, a group of related human developmental overgrowth syndromes, including the Weaver, Weaver-like, Cohen-Gibson, and Overgrowth and Intellectual Disability syndromes, appears to be caused by heterozygous mutations in *EED*, *EZH2* or *SUZ12* (Burkardt et al., 2019; Cyrus et al., 2019). These syndromes manifest with peripheral overgrowth, as well as neurological defects and intellectual disability. Our finding of a particular sensitivity of DA, Hcrt and Tac2 neurons to loss of PRC2 activity may suggest that behaviours related to these hypothalamic cell types may be important to assess in the human PRC2 syndromes. This applies particularly to the effect of PRC2 on the hypothalamic DA neurons, which are known to repress Growth Hormone release from the pituitary (De Zegher et al., 1993; Van den Berghe et al., 1994a; Van den Berghe et al., 1994b), hence forming a plausible link between DA neuron loss and general overgrowth in the PRC2 syndromes.

## ACKNOWLEDGEMENTS

We are grateful to Jose Dias, Johan Ericson, The Jackson Laboratory mouse stock centre and the ENCODE consortium, for sharing reagents and advice. We thank Nigel Kee and Brian Key for critically reading the manuscript. Carolin Jonsson provided excellent technical assistance.

## MATERIALS AND METHODS

### Mouse stocks

The *Eed^fl/fl^* allele has exons 3-6 flanked by *loxP* sites (Xie et al., 2014), and was obtained from Jackson Laboratory Stock Center (Bar Harbor, Maine, USA; stock number #022727). The *Sox1-Cre* line (Takashima et al., 2007) was provided by Jose Dias and Johan Ericson (Karolinska Institute, Stockholm, Sweden). Stocks were maintained on a *B6:129S1* mixed background. Mice were maintained at Linkoping University animal facility, in accordance with best practices. All mouse procedures were approved by the regional animal ethics committee (Dnr 69-14). The morning of the vaginal plug was set as 0.5 days post coitum i.e., E0.5. Pregnant females were sacrificed, and embryos dissected at E12.5, E13.5, E15.5, E16.5 and E18.5. Primers for genotyping were: Cre1: GCG GTC TGG CAG TAA AAA CTA TC. Cre2: GTG AAA CAG CAT TGC TGT CAC TT. Eed1: GGG ACG TGC TGA CAT TTT CT. Eed2: CTT GGG TGG TTT GGC TAA GA. Male/female primers (Clapcote and Roder, 2005); forward: CTG AAG CTT TTG GCT TTG AG; reverse: CCA CTG CCA AAT TCT TTG G.

### Immunohistochemistry

Mouse embryos were dissected to extricate the brains, which were fixed in 4% PFA at +4°C for 18-36h, and kept in 30% sucrose at 4°C until saturated, upon which they were frozen in OCT Tissue Tek (Sakura Finetek, Alphen aan den Rijn, Netherlands) and stored at −80°C. Cryosections (30 µm) were treated with 4% PFA for 15 min at room temperature, blocked and processed with primary antibodies in PBS with 0.2% Triton-X100 and 4% horse serum overnight at +4°C. Secondary antibodies, conjugated with AMCA, FITC, Rhodamine-RedX, Cy5 (Jackson ImmunoResearch, West Grove, PA, USA), or AFD555, AF568 or AF647 (Thermo Fisher Scientific, Waltham, MA, USA), were used at 1:200. Slides were mounted in Vectashield (Vector Laboratories, Burlingame, CA, USA). DAPI was included in the secondary antibody solution. Primary antibodies: Goat anti-Sox2 (1:250, #SC-17320, Santa Cruz Biotechnology, Santa Cruz, CA, USA). Rabbit anti-H3K27me3 (1:500, #9733, Cell Signaling Technology, Leiden, Netherlands). Rat anti-PH3-Ser28 (1:1,000; Cat.no. ab10543). Rabbit anti-Pax2 (1:100, #ab232460), rabbit anti-TFAP2b (1:100, #ab221094), rabbit anti-Orexin A (Hcrt) (1:1,000, ab6214), rabbit anti-Avp (1:500, #ab213708, Abcam, Cambridge, UK).

### Confocal Imaging and Data Acquisition

Fluorescent images were obtained with Zeiss LSM700 or Zeiss LSM800 confocal microscopes. Confocal stacks were merged using Fiji software (Schindelin et al., 2012). Compilation of images and graphs was done in Adobe Illustrator.

### Quantification of proliferation

On confocal images of sagittal sections, a selection was made along the rim of the hypothalamic tissue, of 400 μm width (segmented line), using ImageJ (Fiji) software. This selection was straightened. A second sub-selection of 1000 μm in length of the straightened tissue was made. The final tissue selection analysed was 400 x 1000 μm.

PH3, DAPI and Sox2 staining were quantified using Fiji software (Schindelin et al., 2012; Schindelin et al., 2015). 3D reconstruction and volume quantification of DAPI and Sox2 signals were achieved using 3DViewer Fiji plugin (Schmid et al., 2010) in the selected regions and considering the anatomical features, excluding non-CNS tissue. Mitotic cells (PH3+) were counted in the selected region (mentioned above). Proliferation analyses are presented as ratios of mitotic cells to DAPI or Sox2 calculated volumes within the selected regions. ImageJ (Fiji) thresholding methods (functions) “Huang” & “Moments” were used to remove background/noise from signal in volumetrics analysis of Sox2 and DAPI, respectively.

### Mitotic analysis

For mitotic analyses, two-tailed Student’s T-test was performed. One (*), two (**) or three (***) asterisks were used to assign statistical significance of p≤0.05, p≤0.01 or p≤0.001 respectively. Microsoft Excel 2010 and GraphPad Prism 8.3.0 were used for statistical analyses, data compilation and graphical representation.

### scRNA-Seq

#### Data generation

E12.5, E13.5, E15.5, E16.5, & E18.5 embryos (control: E12.5, E13.5, E15.5, E16.5 and E18.5; *Eed-cKO*: E13.5, E15.5 and E18.5) embryos were harvested and genotyped. The hypothalamus was dissected out in ice-cold RPMI-1640 medium (cat.no. 11530586, Thermo Fisher Scientific, Waltham, MA, USA). The dissected hypothalamuses were further divided into 2-4 pieces before cell dissociation for single cell isolation using Papain Dissociation System (cat no. LK003150, Worthington Biochemical Corporation, Lakewood, NJ, USA). The tissues were incubated with papain solution at +37°C for 60 min, under slow shaking (80 rpm). Isolated single cells were re-suspended in ice-cold 1xPBS, 0.1% BSA, to a final concentration of 2,500 cells/μl (+/- 10%). Cells were checked for viability using a Bio-Rad TC10 automated cell counter (Bio-Rad, Hercules, CA, USA). Single cells and barcode beads were encapsulated into droplets using Bio-Rad ddSEQ Single-Cell Isolator for cell lysis and barcoding (cat.no. 12004336, Bio-Rad, Hercules, CA, USA). Subsequently, RNA-seq libraries were generated using Illumina SureCell WTA 3’ Library Prep Kit for the ddSEQ System (6 cartridge version, cat.no. 20014280, Illumina, San Diego, CA, USA). Libraries were assessed for quality, using 1µl of undiluted cDNA, on an Agilent Technology 2100 Bioanalyzer, using an Agilent High Sensitivity DNA chip (cat.no. 5067-4626, Agilent, Santa Clara, CA, USA) to determine fragment size and yield. Samples were stored at −20℃ until sequencing. Libraries were normalized and sequenced in pools of four, whereby libraries prepared on the same ddSEQ cartridge were balanced between different library pools to control for batch effects. The libraries were sequenced on an Illumina NextSeq500 system, using NextSeq 500/550 High Output Kit v2.5 (150 Cycles, cat.no. 20024907, Illumina, San Diego, CA, USA), paired-end read type and Single-Index. This platform does not have physically separate sequencing lanes, and thus pooled samples were sequenced over all four lanes of the NextSeq flow cell. The scRNA-seq fastq files were generated using the Illumina Base Space application *FASTQ Generation* v1.0.0.

### Cell count matrix generation and quality control

Cell Unique Molecular Identifier (UMI) counts were generate using the Illumina Base Space application *SureCell RNA Single-Cell* v1.2.0; which uses using *STAR* (v2.5.2b) to align the cDNA reads to the mm10 genome (the second read mate, R2) and *samtools* (v1.3) to append cell barcodes to aligned reads (derived from the first read mate, R1). The generated counts for each sequencing run in each sample were concatenated into a single count matrix; yielding a total of 20,703 cells. For technical quality control, cells with fewer than 600 UMIs, less than 1.2 UMIs/expressed gene, and less than 300 genes expressed were removed. This resulted in 14,121 high quality cells (All-Cells). These had median values of 4,559 UMIs per cell, 2,430 expressed genes per cell, and 1.87 UMIs/gene expressed (Figure S3).

We then performed biological filtering in order to remove cells which do not develop within the hypothalamus. For these steps, we used ∼500,000 reference single cell expression profiles with cell type labels (TaxonomyRank4 labels) from the mouse nervous system (http://mousebrain.org/downloads.html) (Zeisel et al., 2015), in combination with *scmapCluster* from *scmap* v1.4.1, to obtain initial cell type labels. Cells that do not develop within the hypothalamus were filtered, which included vasculature and immuno-related cell types (Vascular Endothelial, Vascular/Leptomeningial, Pericytes, Microglia).

Cells expressing *Foxg1* were also removed, since *Foxg1* expression demarcates the anterior extent of the hypothalamus (Figure 1). This resulted in 11,713 hypothalamic cells (All-Hypo). Cells were then normalized according to counts per ten-thousand (CPT) and log transformed (log2((gene-UMIs/cell-UMIs)*10 000)+1, i.e. log2(CPT+1)). The same biological filtering and quality control procedure was also applied to cells generated at the earlier embryonic time points (Figure S10).

### Dimensionality Reduction

The main goal of our dimensionality reduction procedure was to capture a joint space of the control and Eed-cKO cells that would allow direct comparison of cellular diversity. Toward this end, we utilised *Principal Components Analysis* (*PCA*) for initial dimensionality reduction. PCA requires the input data to be mean-centred and variance-normalised per gene. In order to satisfy these requirements, we mean-centred and scaled to unit variance our normalised gene expression matrix using the *scale* function is *scikit-learn v0.21.3* prior to any PCA transformation. Importantly, we then derived the PCA transformation on the control cells alone, and then applied the same transformation on the *Eed-cKO* cells. This ensured that our dimensionality reduction captured the main dimensions of variation in the control cells and not sources of variation due to the *Eed-cKO* treatment; hence allowing a direct comparison between these two conditions without complicated batch correction techniques. The PCA spaces of the two sets of cells were then concatenated, and *Uniform Manifold Approximation and Projection* (*UMAP*) was performed on this joint space using *umap-learn* v0.3.8. The genes used for each PCA prior to UMAP embedding are provided in Table S1; these were selected depending on the set of cells analysed, as detailed below.

### Gene Selection

To prioritise the biological variation, we were interested in – primarily neurotransmitter, neuropeptidergic and DArgic diversity – we curated lists of marker genes for such populations. To counteract technical noise from utilising few genes we selected additional genes based on their correlation with these marker genes (as detailed for the specific analyses below). This approach was inspired by the method Garnett, which performs similar analyses for cell type prediction (Pliner et al., 2019). As evidenced by our resulting clustering and UMAP embeddings, this approach successfully allowed us to analyse hypothalamic cellular diversity in our control and *Eed-cKO* mice.

### E18.5 All-Hypo Cells Analysis

To investigate the effect of *Eed-cKO* on broad hypothalamic cell types, we compiled a list of marker genes for neurons (GABAergic, glutamatergic) and glia (oligodendrocytes, astrocytes, tanycytes, ependymal cells). These are provided in Table S1. We subsequently labelled cells according to their type based on which marker gene they expressed the most highly. Cells not found to express any of these marker genes but expressing NP-DA markers were labelled as ‘other-neuron’. Cells not expressing any differentiated cell type marker were considered un-labelled.

We then selected additional genes which would maximally separate our cell type labels using *normalised mutual information* (NMI, *scikit-learn* v0.21.3), a measure of correlation between discretely labelled data. To obtain discrete labels for gene expression, we classified genes as ‘on’ (counts>0), or ‘off’ (counts=0). We then selected the top 300 genes with the highest NMI (see Table S1) between the binary expression of that gene and the cell type labels. Using these marker genes, the dimensionality reduction procedure was performed as detailed above; with the top 100 principal components using 50 nearest neighbours, a minimum distance of 0.3, and the correlation distance metric as input to UMAP.

### E18.5 NP-DA Cells Analysis

Cells were considered NP-DA if they expressed any of 32 neuropeptidergic or DA-ergic marker genes compiled from the literature (Table S1). 8,219 hypothalamic NP-DA cells were isolated from the rest of the cells prior to application of dimensionality reduction (detailed above) based on the 32 marker genes, along with the top 90 genes that had the highest Pearson correlation coefficient for at least one of the marker genes. The UMAP was performed based on the top 60 principal components using 10 nearest neighbours, a minimum distance of 0.01, a spread of 10, and the Manhattan distance metric.

### E18.5 NP-DA Cell Clustering and Differential Gene Expression

Within the UMAP space containing both the control and *Eed-cKO* NP-DA cells, a nearest neighbour graph was constructed using the kneighbours_graph function in sklearn, with 4 neighbours and the Euclidean distance measure. The cells were subsequently clustered using the Lieden algorithm (*liedenalg v0.8.2*) with a resolution of 2. This resulted in 149 clusters. To derive the meaning behind the clustering, we used differential gene expression to label clusters according to the combination of NP-DA markers they upregulated. Significantly differentially expressed genes were determined for each cluster using independent t-tests *(scipy v1.4.1*), with a one-versus-rest mode of comparison (Bonferroni p-value < 0.05). We found this method of calling differential expression yielded the most reasonable number of differentially upregulated genes in order to distinguish each cluster, as opposed to Student’s t-test which yielded many clusters with no NP-DA markers DE, and Wilcoxon rank-sum tests which yielded many clusters with numerous NP-DA markers DE (data not shown).

In cases where no NP-DA marker was found to particularly distinguish that cluster, clusters were labelled with the NP-DA markers with the highest proportion of expression in that cluster along with the best distinguishing gene in the extended list of genes used to derive the NP-DA UMAP (Table S1). Clusters were subsequently merged if they were found to upregulate the same combination of NP-DA marker genes and were neighbours in the UMAP space, or the cluster was not found to upregulate any marker but was spatially close in the UMAP space to another cluster and showed clear visual expression of the same marker gene.

This manual curation resulted in 116 hypothalamic NP-DA clusters. The aforementioned method for calling differentially expressed genes was subsequently re-run to ensure the markers remained DE in the final clusters; and to call differentially expressed genes within clusters between *Eed-cKO* and control cells (Table S5).

The heatmaps were constructed by taking the average expression of each gene in each NP-DA cluster segmented by control and *Eed-cKO*. The average expression of each gene across the NP-DA clusters was scaled between 0 and 1, to thereby visualise the expression in each cluster relative to the total across the clusters. Scaling was performed to enable direct comparison between the control and *Eed-cKO* heatmaps. These visualisations were constructed using *seaborn* v0.11.0.

### E18.5 Broad Differential Expression and Gene Set Enrichment

Differential expression of control versus *Eed-cKO cells* - regardless of cluster - was performed using DEsingle (Miao et al., 2018). This resolved genes into different types of differential expression; differential abundance of cells expressing the gene (DEa); differential magnitude of gene expression (DEm); or general differential expression (DEg), where there is a change in both the abundance of cells expressing the gene and the magnitude with which that gene is expressed. This differential expression was performed for the All-Hypo cells and the NP-DA cells separately (Table S2).

To investigate if the grouping of the genes into the types of differential expression had a biological significance, we performed gene set enrichment using the *Enrichr* method, grouping the genes into up- or down-regulated genes for each of the DEa, DEm, and DEg genes using gseapy v0.9.16 (Figure S4). We tested for enrichment using the *GO_Molecular_Function_2018* and the *ENCODE_Histone_Modifications_2015* gene sets, thus giving us insight into the molecular function and the epigenetic modifications of these genes across tissues (adjusted p≤0.5; Tables S3 and S4).

When visualising these results, we grouped terms based on similarity (Figure S4). Terms were grouped as “TF activity/DNA binding” if they mentioned “binding” or “transcription”. Terms were grouped as “H3K27me3”, “H3K36me3” or “H3K4me3” if either of these epigenetic marks were mentioned in the term name; all remaining terms were grouped as ‘Other’. These groupings were based on a manual inspection of the enriched terms which revealed an over-representation of terms related to the aforementioned groupings.

Violin plot visualisations of the DE genes only included 5% of zeros for each grouping of cells, to prevent most of the density falling at zero. In the construction of the ‘Transcription Factor’ violin plot in Figures 1 and S6, we display the top upregulated genes in the *Eed-cKO* that appear in the GO term *DNA-binding transcription factor activity* (GO:0003700). The violin plots were constructed using *seaborn* v0.11.0.

### Early Timepoint Cell Labelling

After quality control, as previously described, we subsequently labelled cells (control: E12.5, E13.5, E15.5, E16.5, & E18.5; *Eed-cKO*: E13.5, E15.5, E18.5) based on expression of known marker genes for glutamatergic, GABAergic, glial, and ventricular zone and subventricular zone cell types; the full cell type and marker list can be seen in Table S8. Due to varying specificity of markers for their respective cell types, we marked cells as belonging to a particular group in a hierarchical manner to alleviate this problem; whereby least specific markers were used to mark cells first, and most specific markers to mark cells last such that prior labelling’s was overridden. The order this was performed is also shown in Table S8, where groups labelled first to last are listed from the top row of the table to the bottom row of the table.

A gene was considered ‘on’ in a particular cell if that cell expressed the gene in the upper 40^th^ percentile of the non-zero gene expression range of that gene. Labelling of cells was performed based on this binarization (‘on’/’off’) transformation of the gene expression in each cell. In the labelling process, we also considered negative gene expression indicators for particular cell types, and also ‘or’ and ‘and’ logic, where cells could be considered of a certain type if they expressed/did not express indicated marker genes in part or in combination. The exact logical notation used in each case is provided in Table S8.

As a further contingency to prevent labelling cells incorrectly and to reduce the number of unlabelled cells, we then trained a logistic regression classifier (as implemented in sklearn v0.22.2) using all the genes present in Table S8 as well as all mouse transcription factors (as listed in GO term GO:0003700). To train the classifier on relevant genes with this set, we selected the top 300 genes which when considering their binarized expression (expression > 0) that had the highest normalised mutual information score with a randomly selected 80% of cells labelled using marker genes as a training set. We then trained the logistic regression classifier on this training set using the L1 penalty and liblinear solver, and then predicted the labels for the remaining 20% of binary labelled cells and the unlabelled cells. If the probability of the most likely predicted label for a given cell was smaller than 0.5, the cell was labelled ‘unknown’. The confusion matrix for the left out 20% of cells are presented in Table S8, showing the labelling approach taken was effective in labelling the numerous kinds of broad cell types.

### Early Timepoint Dimensionality Reduction

We performed dimensionality reduction as described above; except the gene set used was the top 300 most highly informative genes used for training the logistic regression classifier for the early timepoint cell labelling. The UMAP was then performed by creating a nearest-neighbours graph using scanpy v1.6.1 with 10 neighbours and the top 40 principal components, followed by scanpy’s UMAP function applied with default settings. Notably, for the control cells we sampled a much greater range of time points (E12.5, E13.5, E15.5, E16.5, & E18.5) than *Eed-cKO* (E13.5, E15.5, and E18.5) in order to create a joint space more reflective of the developing hypothalamus. For all subsequent analyses comparing control with *Eed-cKO*, we compare only the overlapping time points (E13.5, 15.5, and E18.5) to prevent biological confounding from the additional time points in the control condition.

### Early Timepoint Identification of *Eed-cKO* Affected Genes in Lhx9 and Glut/GABA cells

We first called differentially expressed genes between control & *Eed-cKO* within the groups of Lhx9 and Glut/GABA cells using a pseudosampling approach followed by a t-test to call DE genes. Namely, we created 20 random samples of approximately equal numbers of cells without replacement within each group/condition combination. The gene expression of these pseudosampled groups were then averaged, and DE genes were called between between *Eed-cKO* and control using a t-test (scanpy) with each pseudosample as an observation. Genes with a false discovery rate adjusted p-value <.05 were taken as significant (Table S9).

The pseudosampling approach was then repeated to compare *Eed-cKO* cells from Lhx9 and *Glut/GABA* cells against all other *Eed-cKO* cells (Table S10). To identify the *Eed-cKO* genes that were specifically upregulated in the *Eed-cKO* Lhx9 and/or Glut/GABA cells, we then took the overlap between the upregulated genes from comparing Lhx9 and Glut/GABA *Eed-cKO* versus Lhx9 and Glut/GABA control cells, and the DE genes from comparing Lhx9-/Glut/GABA *Eed-cKO* cells versus all other *Eed-cKO* cells (Table S11).

## Supplemental Figures and Tables Legends

**Supplemental Figure 1.**
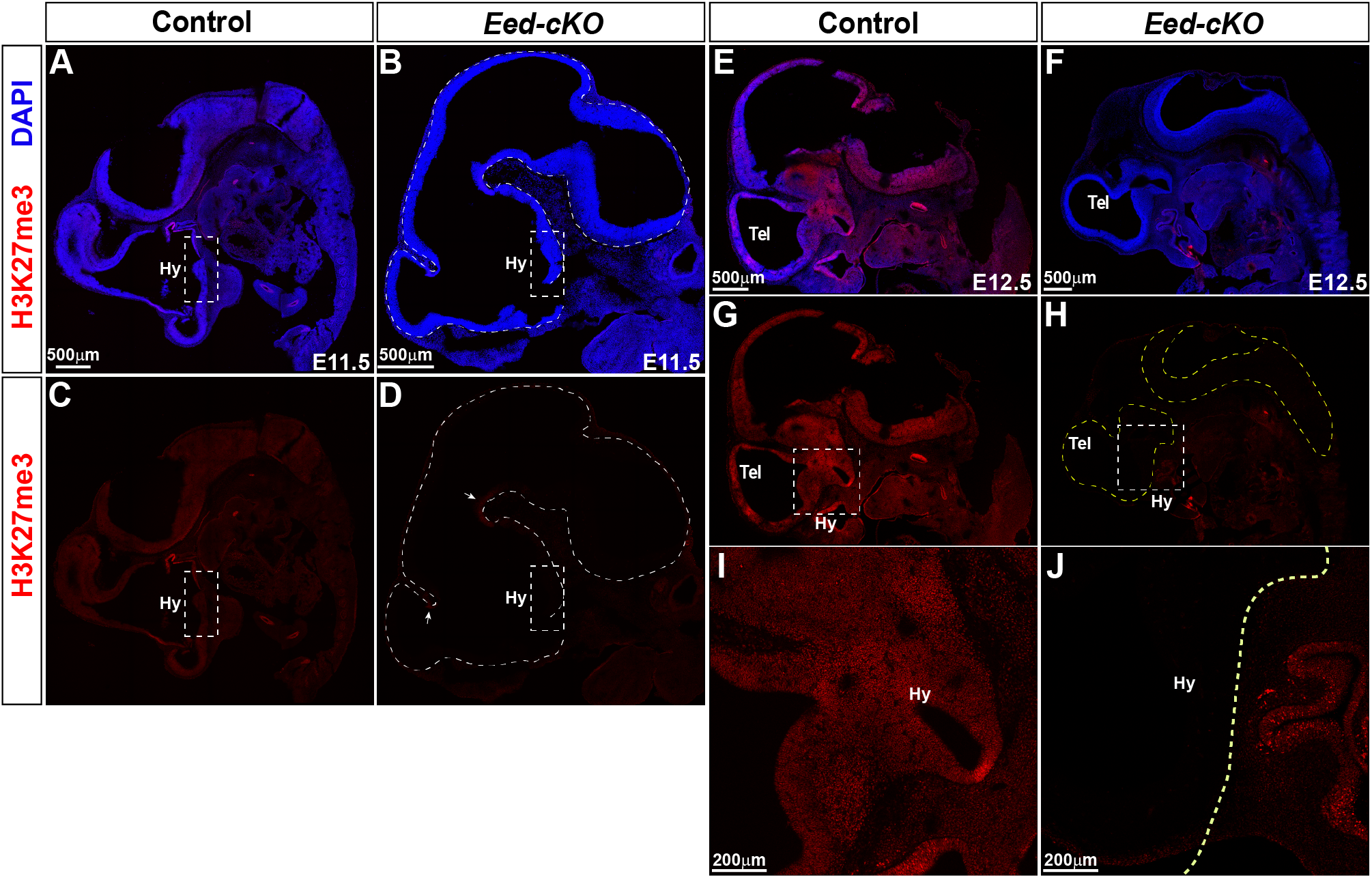
*Eed-cKO* mutants display loss of H3K27me3. (A-D) Sagittal sections of E11.5 control and *Eed-cKO* embryos, stained with DAPI and immunostained for H3K27me3. White dashed line in (D) delineates the CNS in the *Eed-cKO*. In control, H3K27me3 staining is observed throughout the CNS. In *Eed-cKO*, staining is below detection in most parts of the CNS, including in the hypothalamus. A weak signal is apparent in the *Eed-cKO* mutants at the boundary between Tel- and Diencephalon, as well as at the boundary between the Diencephalon and Mid-brain (arrows in D). (E-J) Sagittal sections of E12.5 control and *Eed-cKO* embryos, stained with DAPI and H3K27me3. White, dashed square insets in (G-H) indicate hypothalamic regions in control and *Eed-cKO* magnified in (I) and (J) respectively. Yellow dashed line in H and J delineates the CNS, including the hypothalamic region. In control, H3K27me3 staining is observed throughout the CNS. In *Eed-cKO*, staining is below detection in all parts of the CNS, including in the hypothalamus. Scale bar; 500 µm in (A-H), 200 µm in (I, J).

**Supplemental Figure 2.**
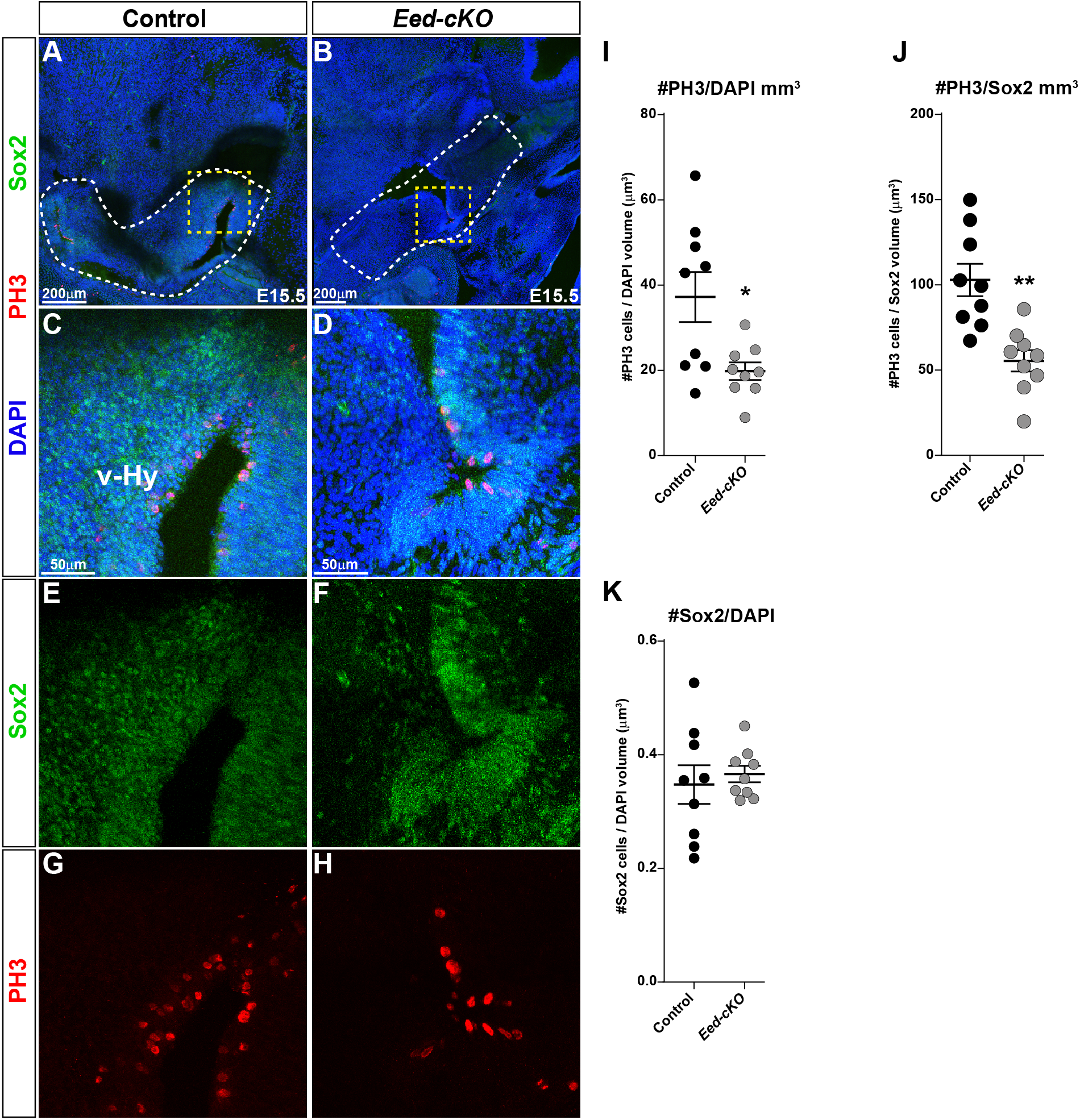
*Eed-cKO* mutants display reduced proliferation in the hypothalamus. (A-H) Staining for Sox2, PH3 and DAPI in sagittal sections of E15.5 embryos, in control and *Eed-cKO*. (A-B) White dashed lines delineate the hypothalamic region used for proliferation analysis. Dashed square insets delineate regions magnified in (C-H). (I, J) Quantification of proliferation in hypothalamic tissue of control vs *Eed-cKO*, at E15.5, plotted as PH3+ cells per mm^3^ of DAPI and Sox2 signals respectively, shows reduced proliferation in *Eed-cKO*. (K) Ratios of Sox2/DAPI signal did not change between hypothalamic tissues of control and *Eed-cKO* embryos at E15.5. Student’s t-test; mean +/- SEM; n= 3 embryos per genotype, n=9 sections; 3 sections per embryo, per genotype. Scale bar; 200 µm in (A, B), 50 µm in (C-H). For source data, see Table S7.

**Supplemental Figure 3.**
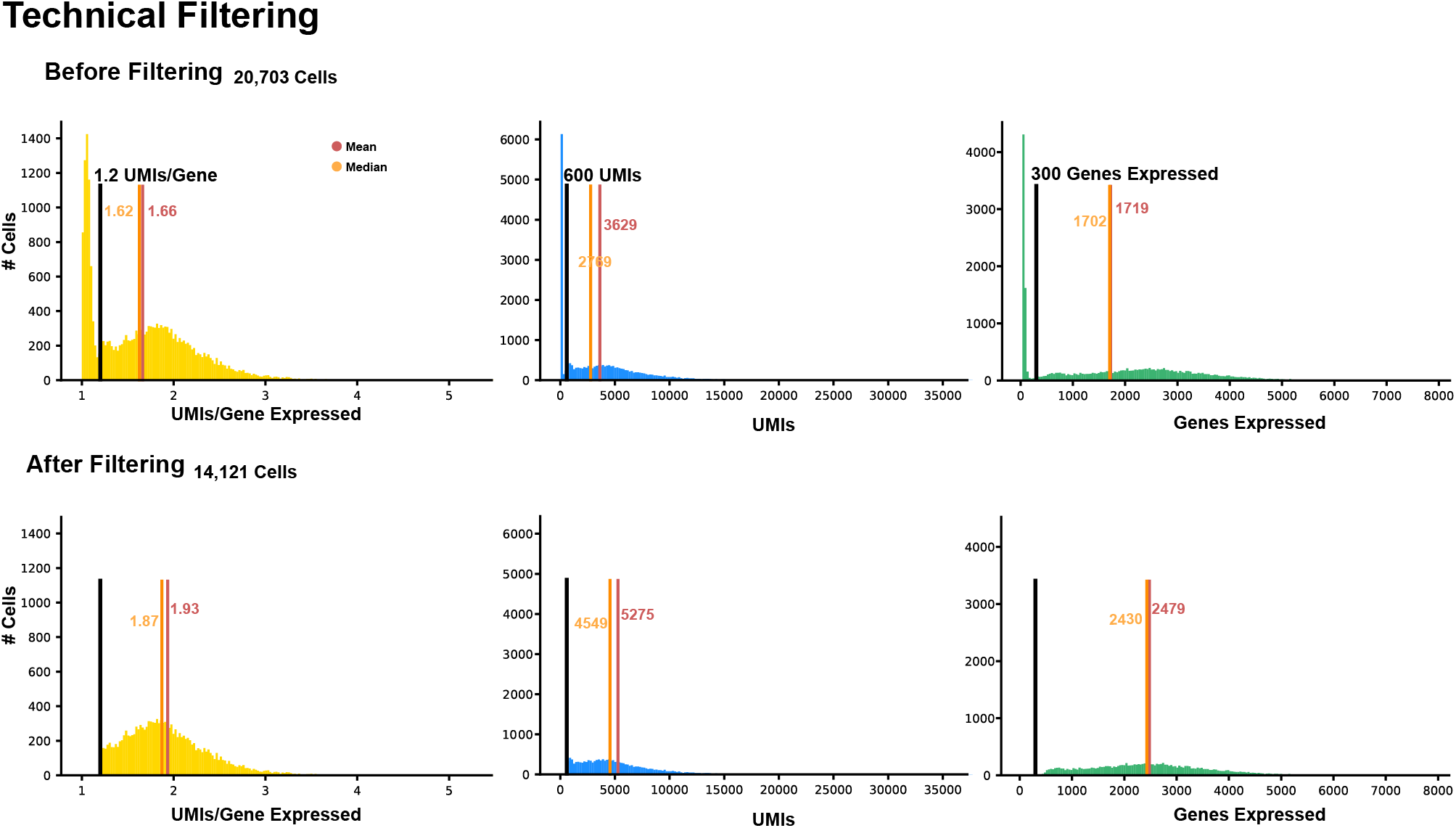
Technical filtering of scRNA-seq data. Technical filtering at E18.5 depicts histograms of the Unique Molecular Identifiers (UMIs) expressed per gene, the total UMIs, and the genes expressed across the cells both before (top) and after (bottom) quality control, respectively. The black, red, and orange vertical lines indicate the cut offs used for quality control, the mean for each quality metric, and the median for each quality metric, respectively.

**Supplemental Figure 4.**
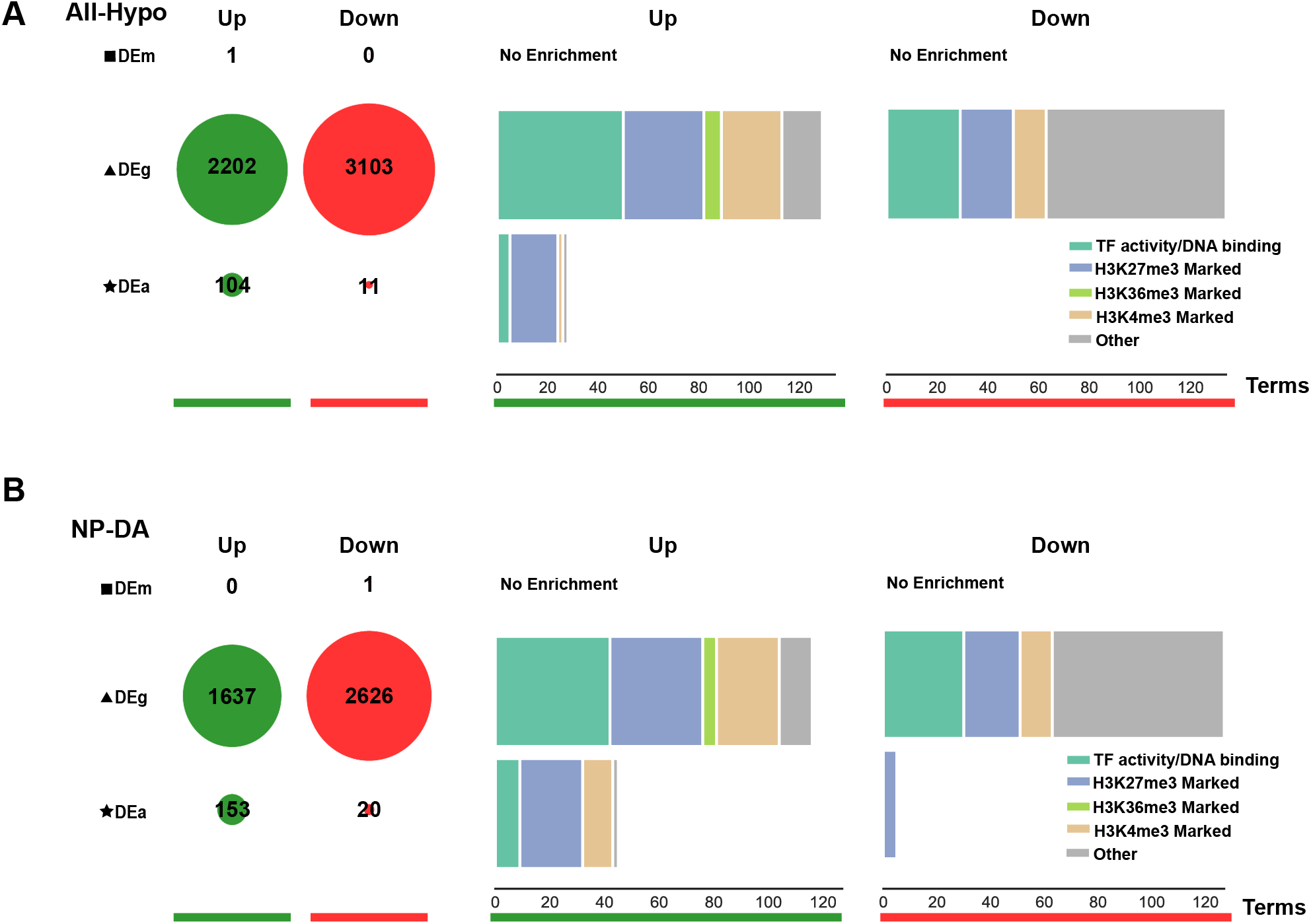
GO analysis of scRNA-seq data. (A-B) GO analysis of All-Hypo (A) and NP-DA cells (B). The first panel (left) shows the total number of genes up (green) and down (red) for each type of differential expression (DEa, DEg, and DEm) in the *Eed-cKO* (adjusted p-value *<0.05).* The right panel depicts stacked bar charts of the total number of gene sets (“Terms”) grouped as “TF activity/DNA binding” if they mention “binding” or “transcription” in the term name, or “H3K27me3”, “H3K36me3” or “H3K4me3” if these epigenetic marks were mentioned in the term name; the remaining terms are grouped as “Other”. The epigenetic terms indicate genes marked by the respective modification in a biological tissue sample in the *ENCyclopaedia of DNA Elements* (ENCODE.

**Supplemental Figure 5.**
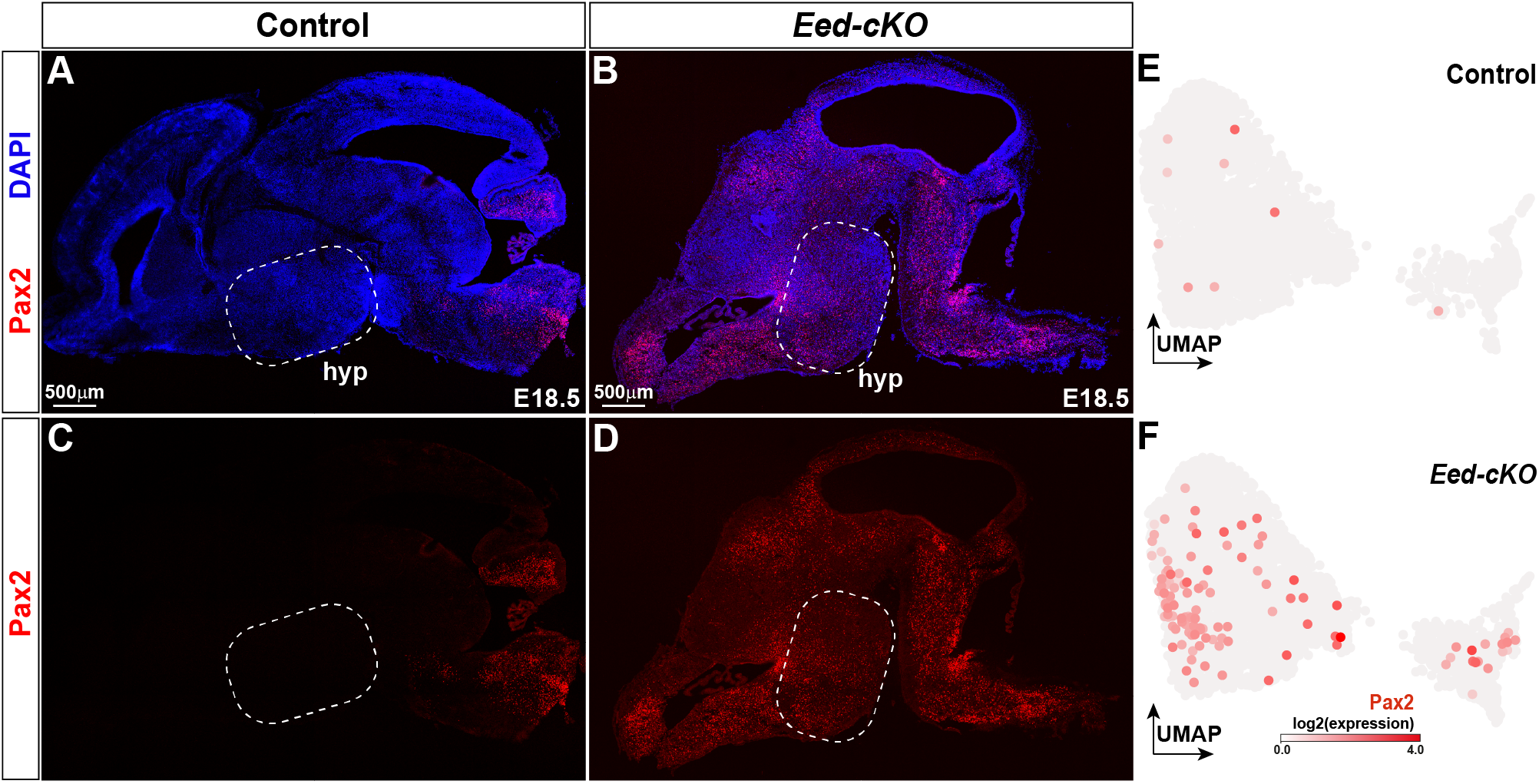
*Eed-cKO* mutants display ectopic Pax2 expression in the hypothalamus. (A-D) Staining for Pax2 and DAPI in sagittal sections of E18.5 brains, in control and *Eed-cKO*. In control, Pax2 is expressed in the mid- and hindbrain. In contrast, *Eed-cKO* mutants display ectopic Pax2 expression in all anterior regions of the brain, including the hypothalamus. (E-F) UMAP embedding of E18.5 hypothalamic scRNA-seq data for the All-Hypo cells, with each cell coloured according to the expression levels of Pax2, in control and *Eed-cKO*. Scale bar; 500 µm in (A-D).

**Supplemental Figure 6.**
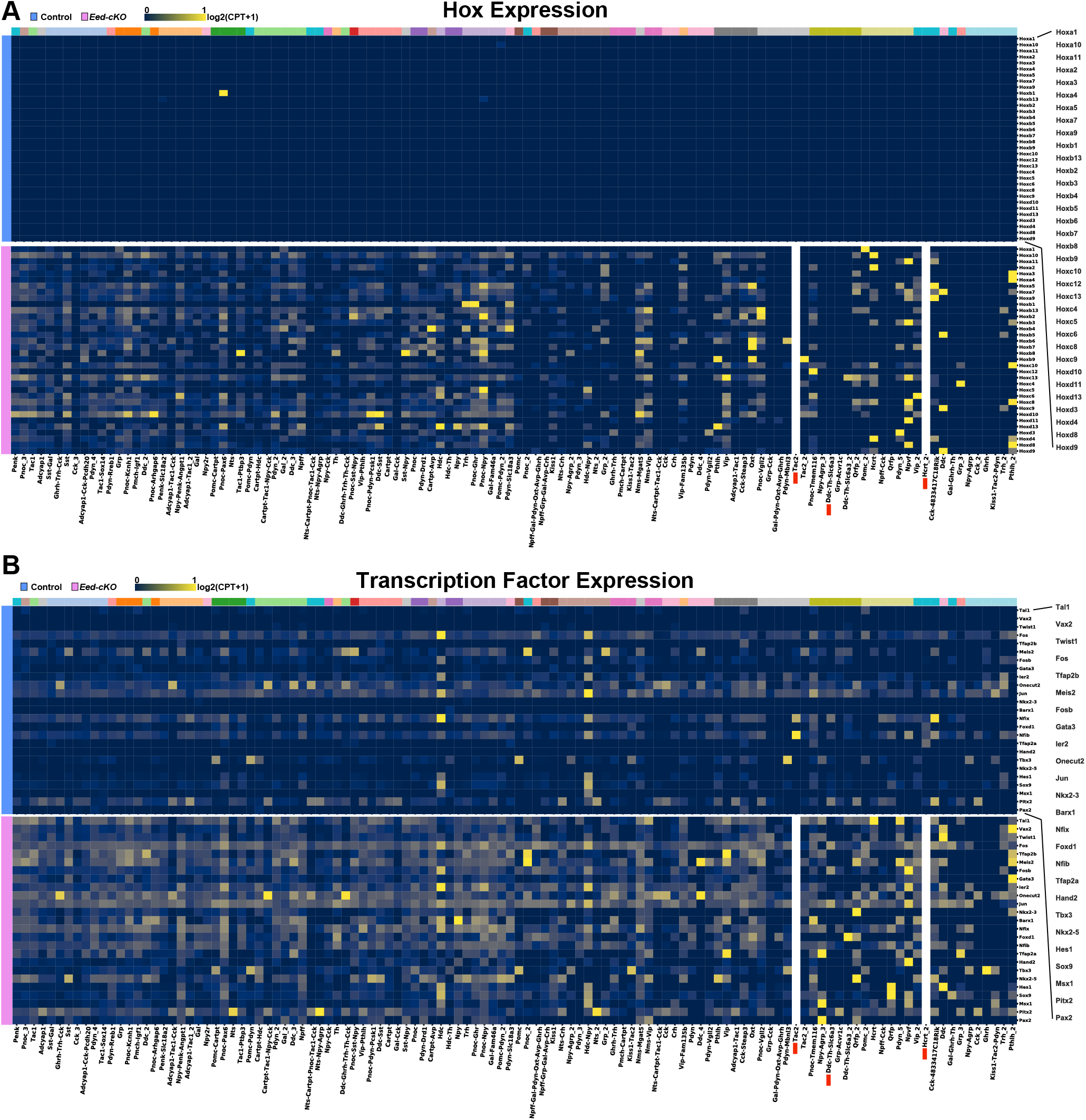
*Eed-cKO* mutants display Hox and posterior TF expression in the hypothalamus. (A-B) Heatmap displaying average gene expression of the Hox and posterior TF genes in each UMAP cell cluster, in control (blue) and *Eed-cKO* (pink). Gene expression is measured in log2(CPT+1), scaled between 0 and 1 along each row. The *Ddc-Th-Slc6a3*, *Tac2* and *Hcrt* clusters are absent in *Eed-cKO*, and hence a column of 0 expression was depicted.

**Supplemental Figure 7.**
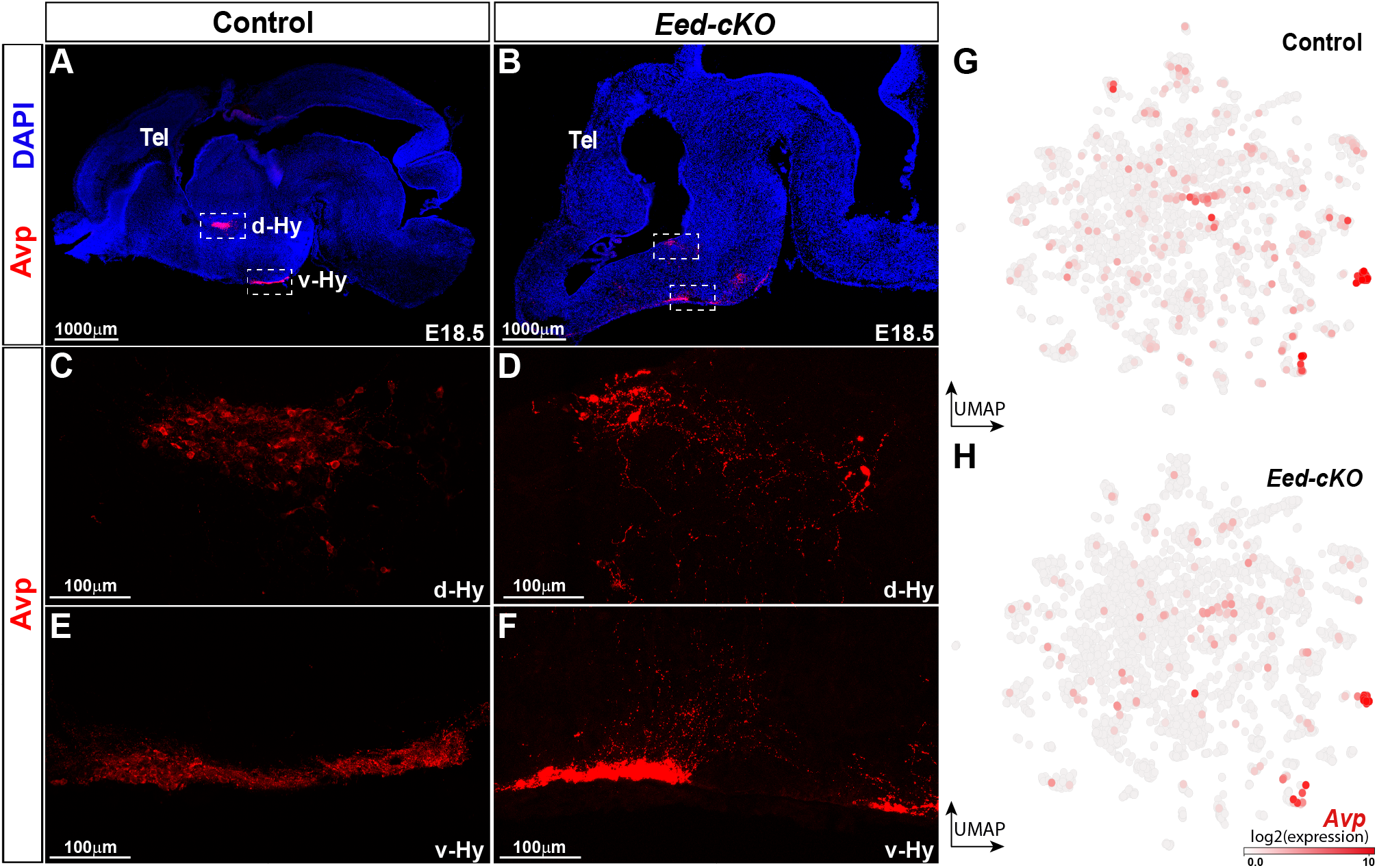
*Eed* is not critical for the generation of Avp neurons. (A-B) Staining for Avp and DAPI in sagittal sections of E18.5 control and *Eed-cKO* mutant brains. Dashed, square insets delineate Avp cells in dorsal and ventral hypothalamic regions. (C-F) Avp+ cells in dorsal (d-Hy) and ventral (v-Hy) regions of hypothalamus in control and *Eed-cKO* embryos. The organization and clustering of v-Hy region Avp cells appear to be disrupted in *Eed-cKO*. (G-H) UMAP embedding of E18.5 hypothalamic scRNA-seq NP-DA cells, with each cell coloured according to the expression level of *Avp* in control and *Eed-cKO*. Scale bar; 1000 µm in (A, B), 100 µm in (C-F).

**Supplemental Figure 8.**
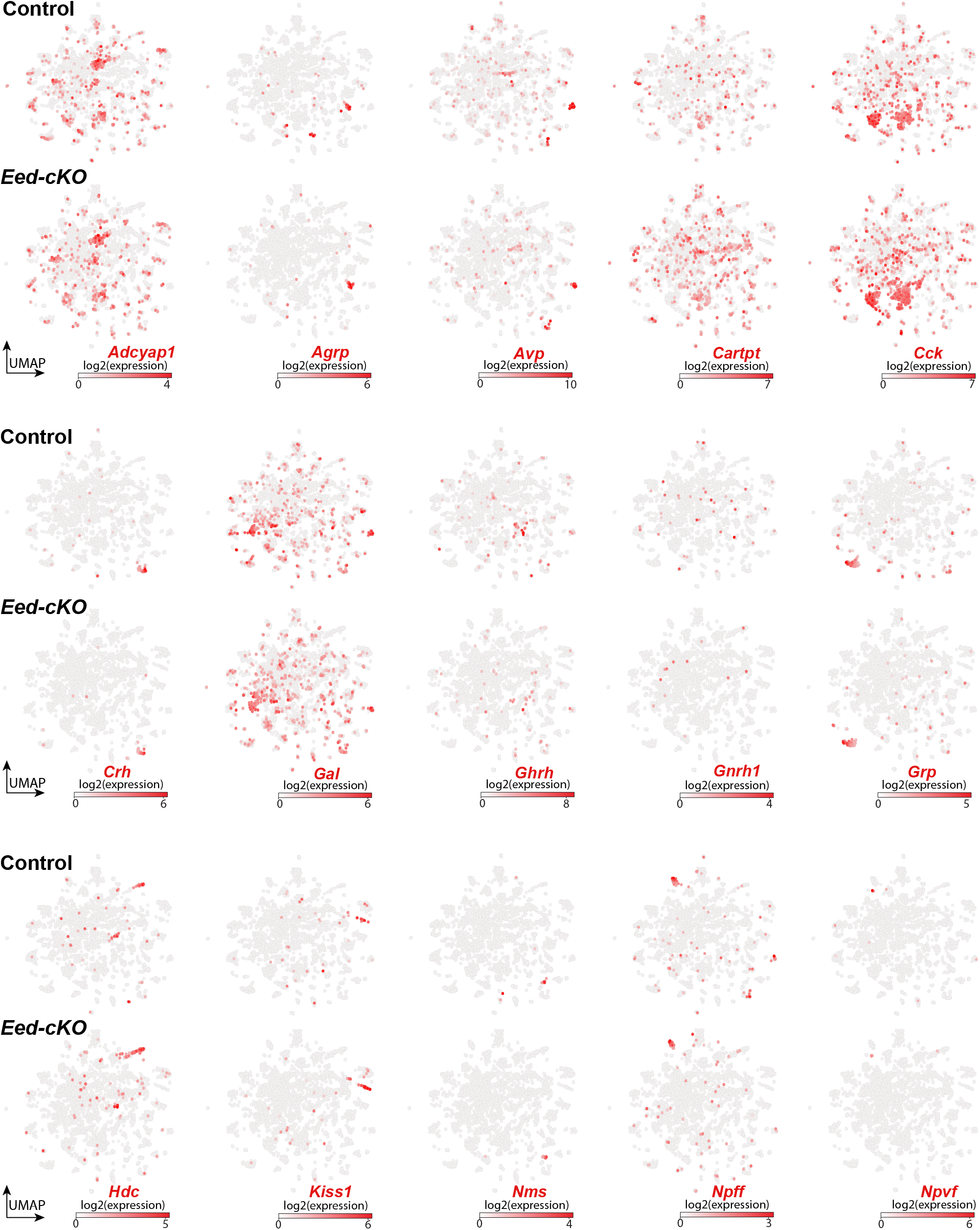
*Eed-cKO* is not necessary for most NP-DA cell fates. UMAP embedding of E18.5 NP-DA cells, based upon 122 DE genes (Table S1), showing expression of neuropeptide genes, in control and *Eed-cKO* cells.

**Supplemental Figure 9.**
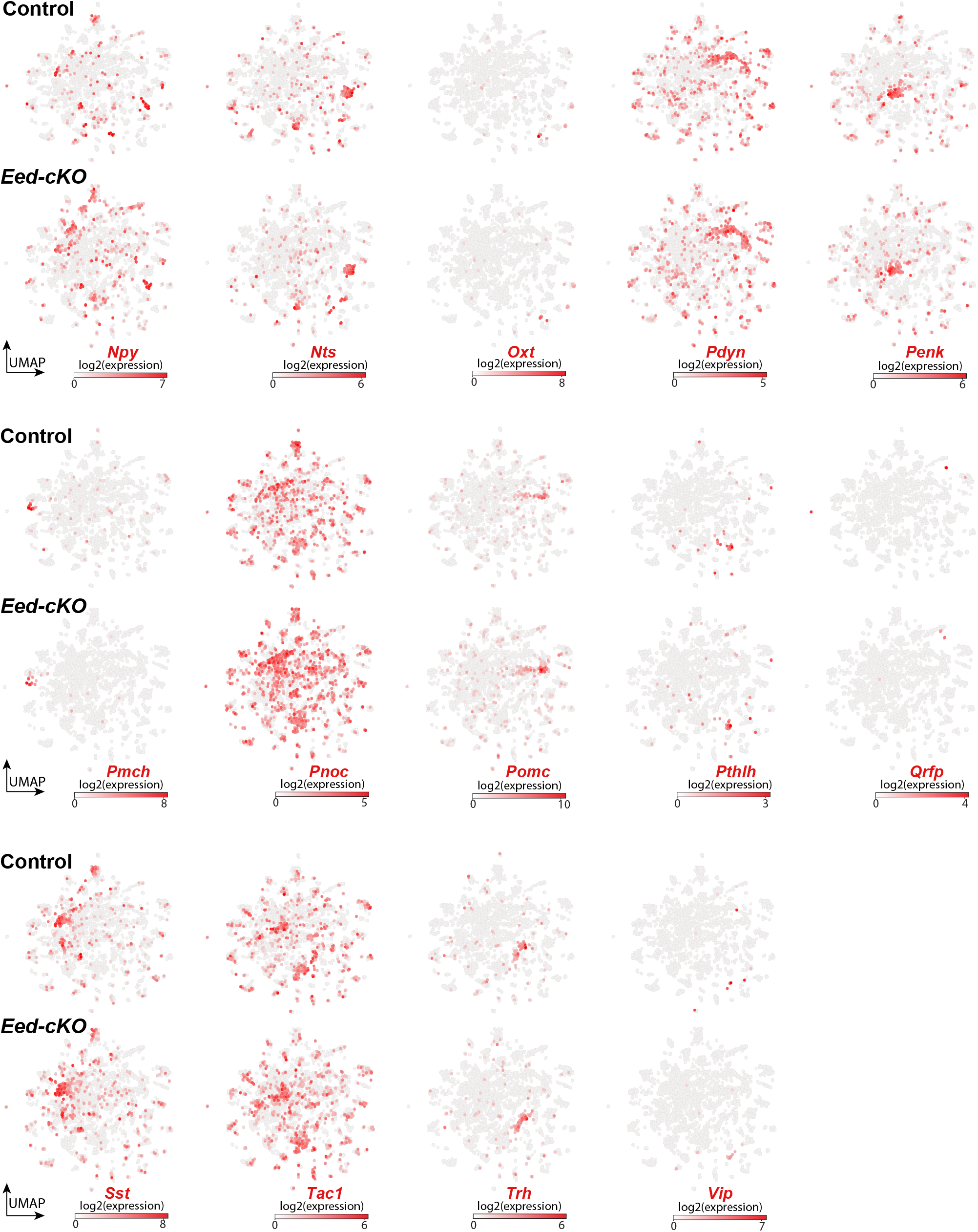
*Eed-cKO* is not necessary for most NP-DA cell fates. UMAP embedding of E18.5 NP-DA cells, based upon 122 DE genes (Table S1), showing expression of neuropeptide genes, in control and *Eed-cKO* cells.

**Supplemental Figure 10.**
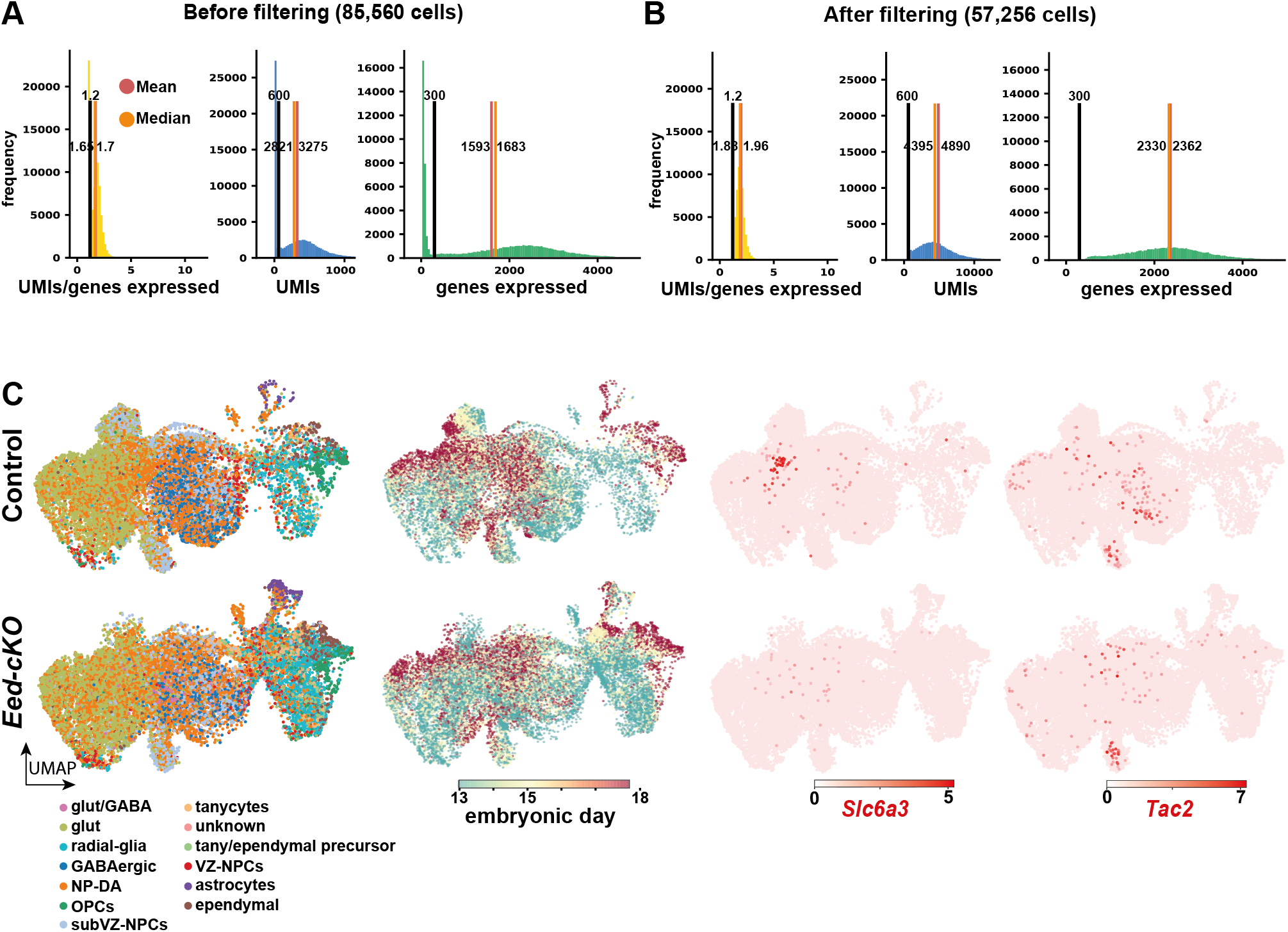
Technical filtering and analysis of E13.5 and E15.5 scRNA-seq data. (A-B) Technical filtering depicts histograms of the Unique Molecular Identifiers (UMIs) expressed per gene, the total UMIs, and the genes expressed across the cells both before and after quality control, respectively. The black, red, and orange vertical lines indicate the cut offs used for quality control, the mean for each quality metric, and the median for each quality metric, respectively. (C) UMAP embedding of E13.5, E15.5 and E18.5 cells, showing that all of the major cell types are generated in the hypothalamus in *Eed-cKO* mutants. However, DA (*Slc6a3*) and a subset of *Tac2* cells are missing.

**Supplemental Table 1.**
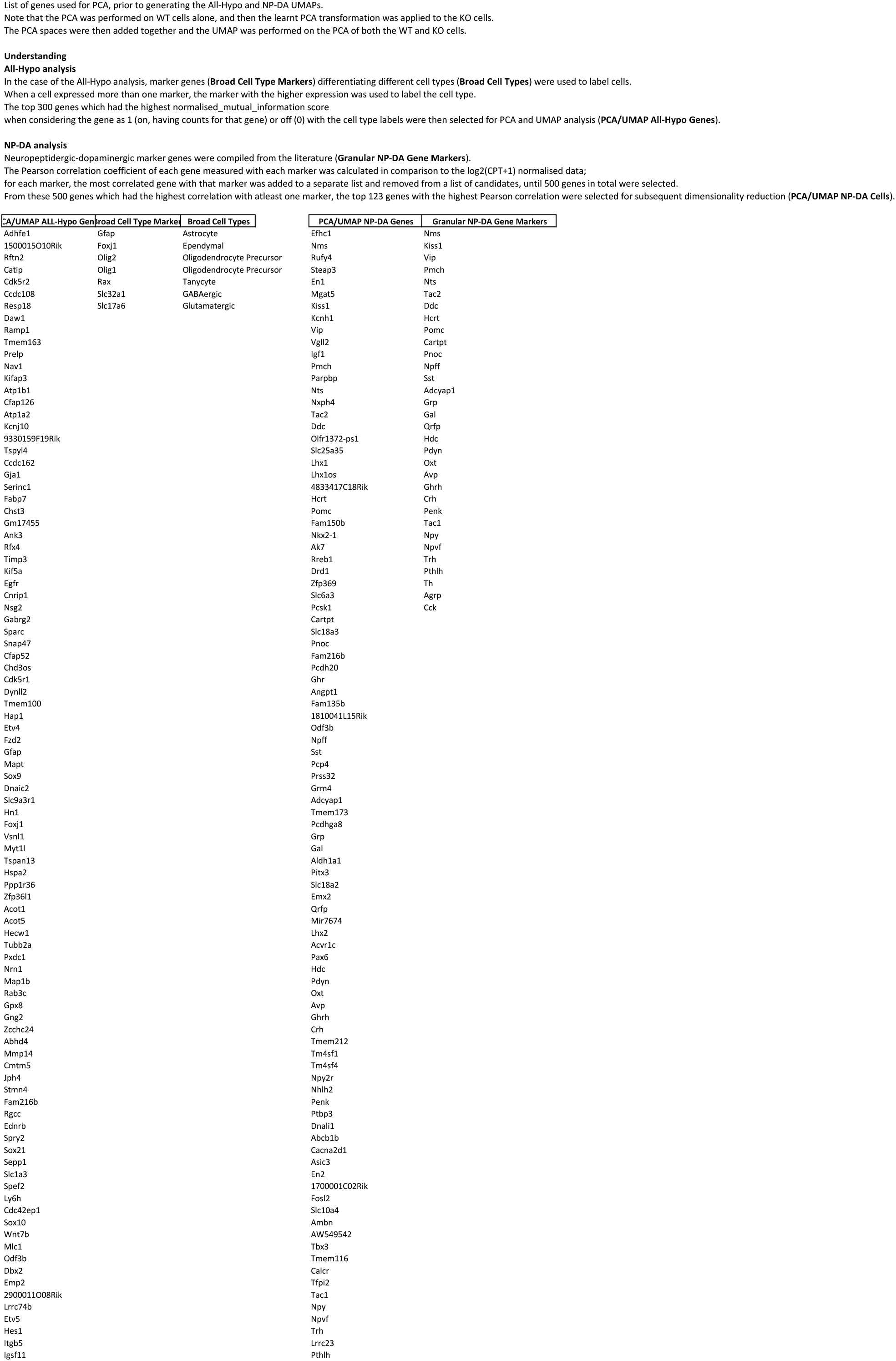
Gene lists for UMAP embedding. Gene lists for UMAP embedding of All-Hypo and NP-DA cells.

**Supplemental Table 2.**
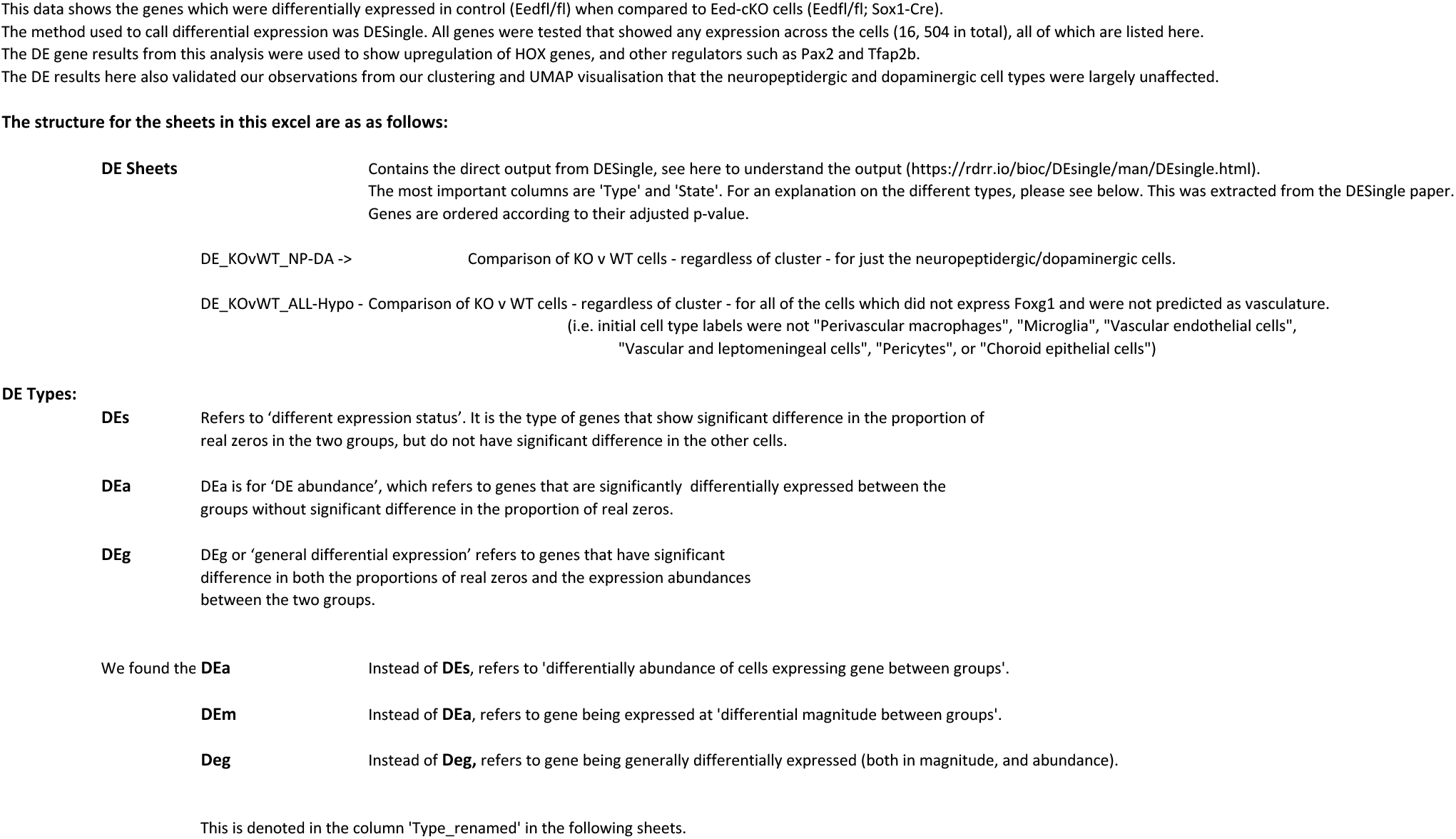
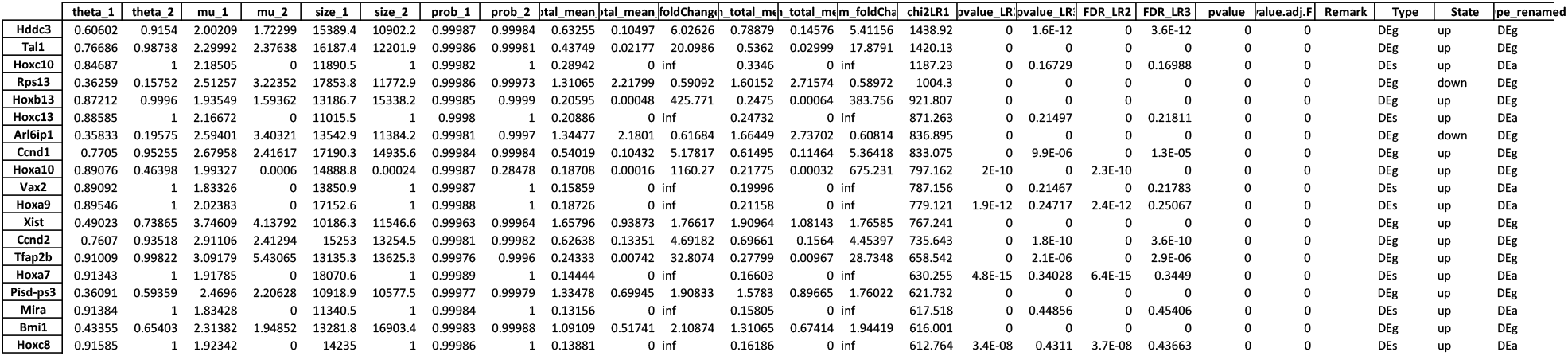
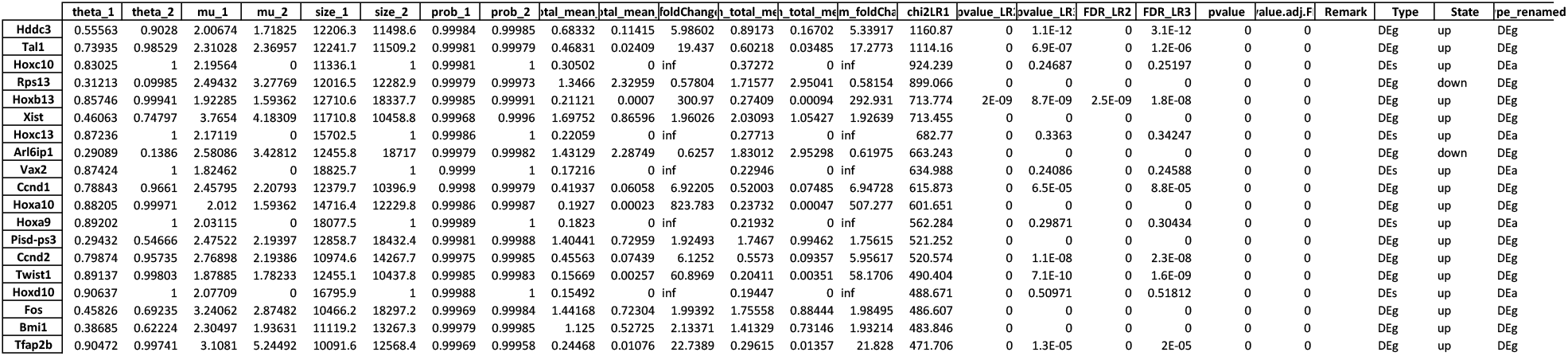
Differential expression in the All-Hypo and NP-DA cells. Differential expression in All-Hypo and NP-DA cells, when comparing single-cell transcriptomic data between control and *Eed-cKO*, at E18.5.

**Supplemental Table 3.**
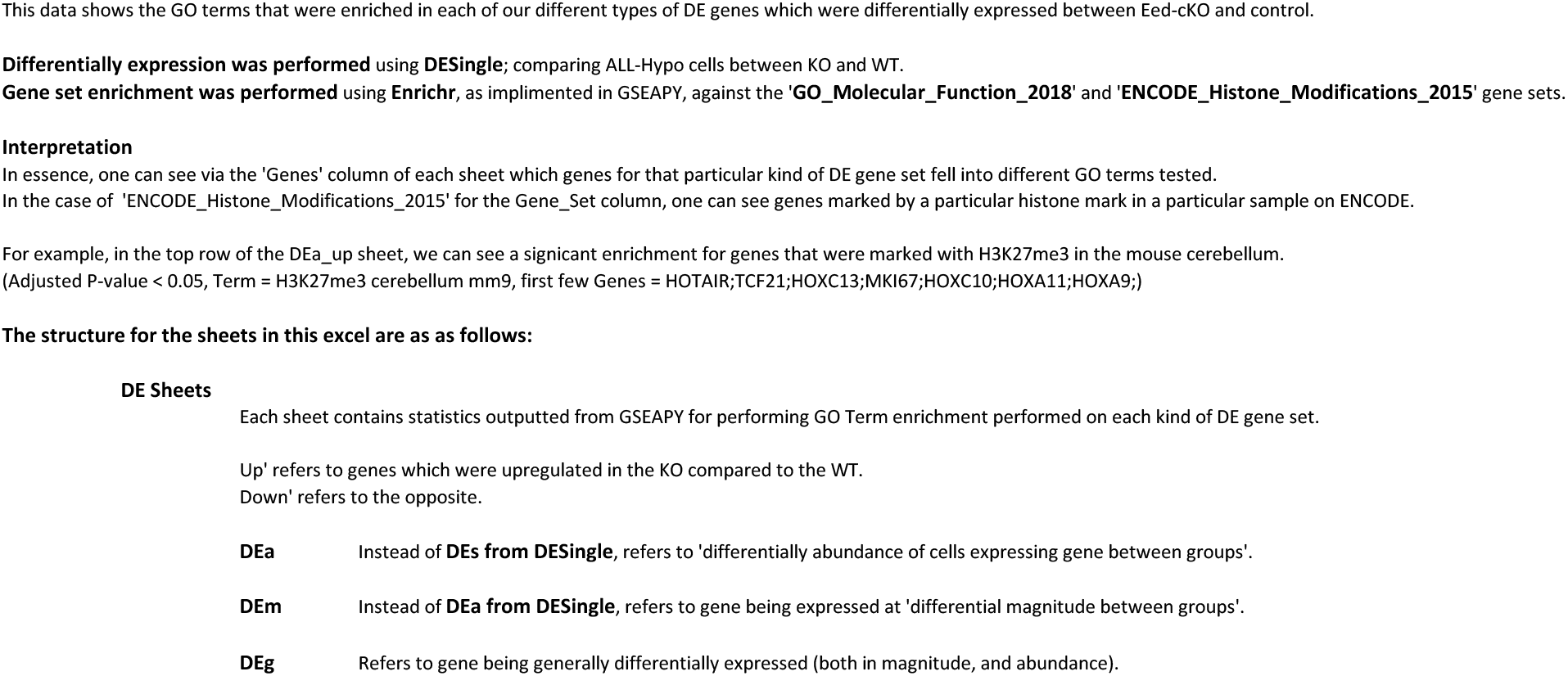
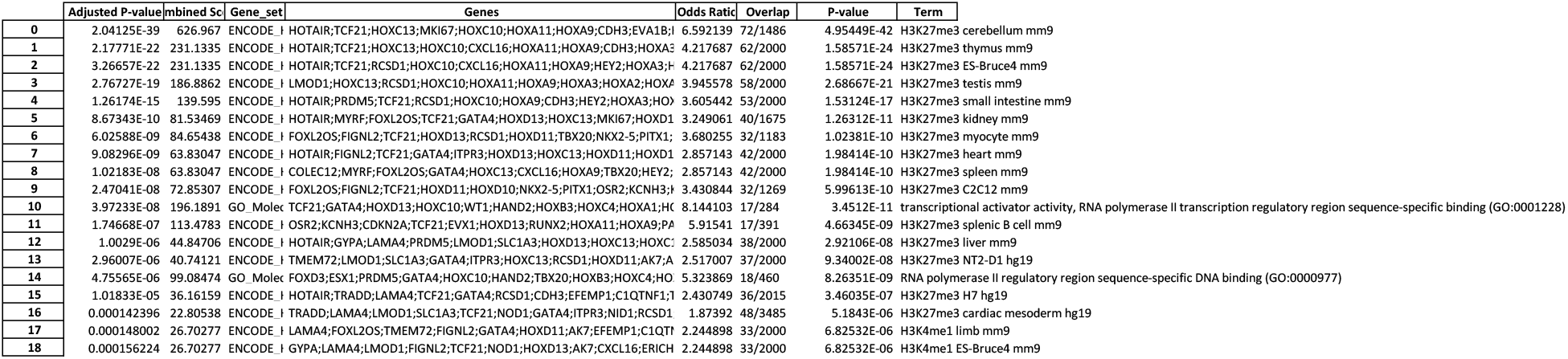
Gene Ontology analysis of All-hypo cells. Gene Ontology analysis of DE genes in All-Hypo cells, at E18.5, showing enrichment of TF activity/DNA binding, H3K27me3 and H3K36me3.

**Supplemental Table 4.**
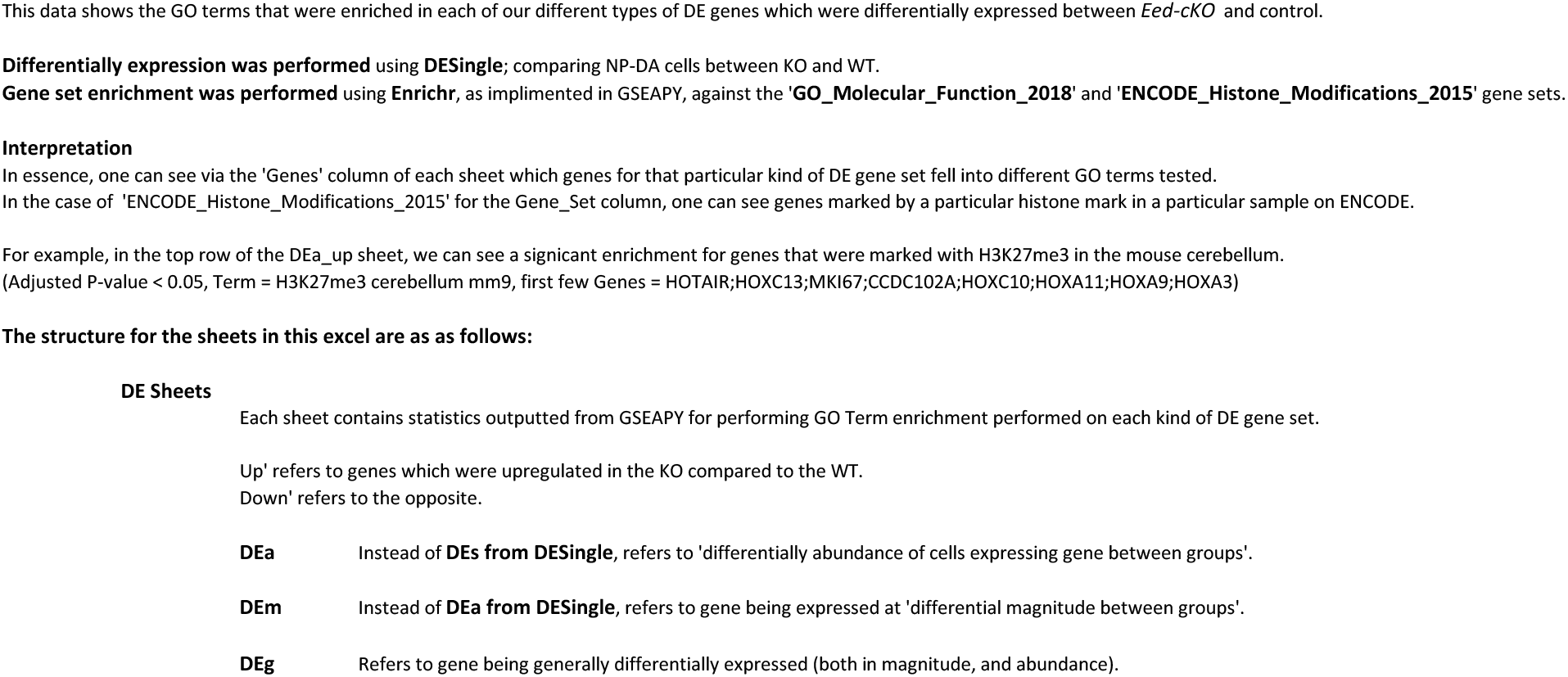
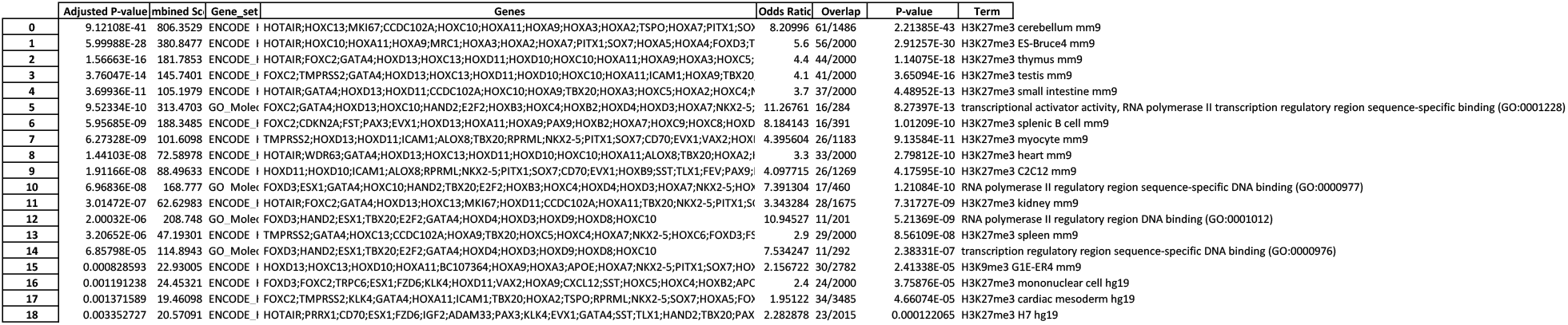
Gene Ontology analysis of NP-DA cells. Gene Ontology analysis of DE genes in NP-DA cells, at E18.5, showing enrichment of TF activity/DNA binding, H3K27me3 and H3K36me3.

**Supplemental Table 5.**
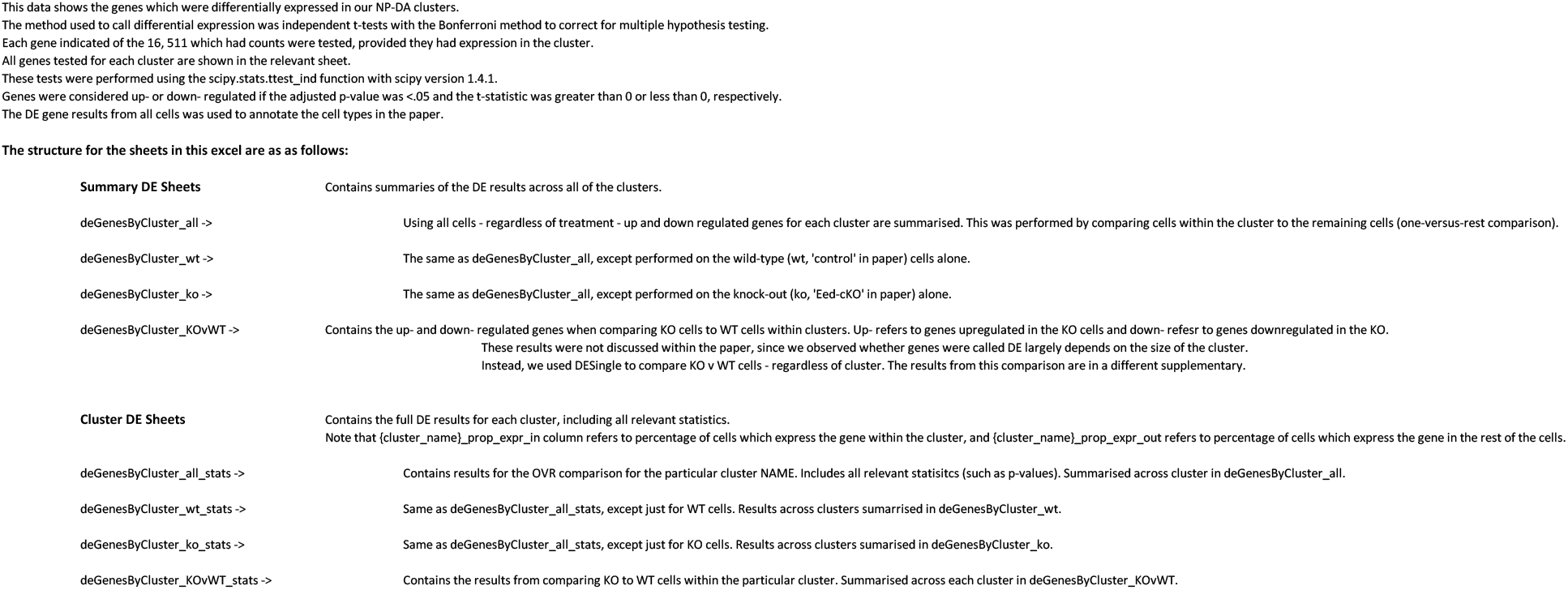
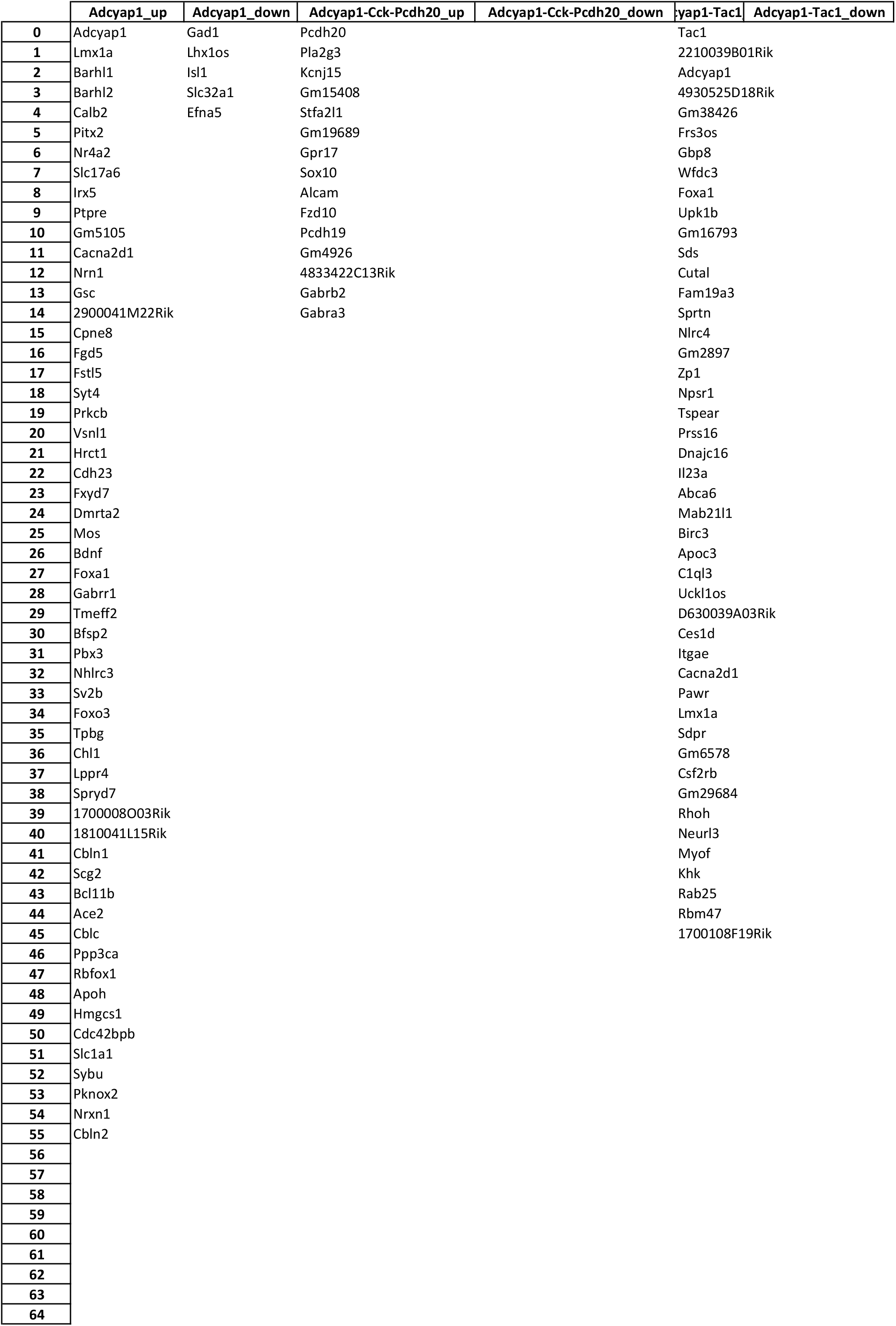
Differential expression in NP-DA cell clusters. Differential expression in NP-DA cell clusters, at E18.5, in control and *Eed-cKO*.

**Supplemental Table 6.** Temporal onset of NP-DA genes and the effects in *Eed-cKO*. Developmental onset of the NP-DA marker genes, based on the Allen Brain Atlas (ABA) (Thompson et al., 2014) and the literature (Diaz et al., 2014; Knoll et al., 2013). Light red depicts weak expression or expression in sub-regions of the hypothalamus.

| <u>E11.5</u> | <u>E13.5</u> | <u>E15.5</u> | <u>E18.5</u> | <u>P4</u> | <u>Transmitters</u> | <u>Ref</u> | <u>DE analysis</u> |
| --- | --- | --- | --- | --- | --- | --- | --- |
|  |  |  |  |  | Adcyap1 | ABA |  |
|  |  |  |  |  | Pomc | ABA | Up DEg |
|  |  |  |  |  | Ddc | ABA | Down DEg |
|  |  |  |  |  | Pmch | ABA | Down DEa |
|  |  |  |  |  | Sst | ABA, Diaz et al. | Up DEa |
|  |  |  |  |  | Th | ABA, Diaz et al. | Down DEg |
|  |  |  |  |  | Trh | ABA, Diaz et al. |  |
|  |  |  |  |  | Tac1 | ABA | Up DEg |
|  |  |  |  |  | Avp | ABA | Down DEa |
|  |  |  |  |  | Cartpt | ABA | Up DEg |
|  |  |  |  |  | Crh | ABA, Diaz et al. | Down DEg |
|  |  |  |  |  | Kiss1 | Knoll et al. |  |
|  |  |  |  |  | Npvf | ABA |  |
|  |  |  |  |  | Npy | ABA | Up DEg |
|  |  |  |  |  | Pdyn | ABA |  |
|  |  |  |  |  | Pnoc | ABA | Up DEg |
|  |  |  |  |  | Grp | ABA | Down DEg |
|  |  |  |  |  | Agrp | ABA |  |
|  |  |  |  |  | Cck | ABA | Up DEg |
|  |  |  |  |  | Gal | ABA | Down DEg |
|  |  |  |  |  | Ghrh | ABA, Diaz et al. | Down DEg |
|  |  |  |  |  | Hcrt | ABA | Down DEg |
|  |  |  |  |  | Nts | ABA |  |
|  |  |  |  |  | Oxt | ABA |  |
|  |  |  |  |  | Penk | ABA |  |
|  |  |  |  |  | Hdc | ABA | Up DEg |
|  |  |  |  |  | Pthlh | ABA |  |
|  |  |  |  |  | Tac2 | ABA |  |
|  |  |  |  |  | Vip | ABA |  |
| ND | ND | ND | ND | ND | Nms |  |  |
| ND | ND | ND | ND | ND | Npff |  |  |
| ND | ND | ND | ND | ND | Prlh |  |  |
| ND | ND | ND | ND | ND | Qrfp |  |  |
| ND | ND | ND | ND | ND | Galp |  |  |

**Supplemental Table 7.** **Source Data for Figure S2** Data and analysis quantifying proliferation in *Eed-cKO* & control.

|  |  |  |  |
| --- | --- | --- | --- |
|  | DAPI-all | volume | 1834281980.00 |
| maxed out value for JAVA-based 32-bit integer | 2.15E+09 |  |  |
| maxed out value for JAVA-based 64-bit integer | 9.22337E+18 |  |  |
| thresholding methods:<br>DAPI: Huang<br>Sox2: Moments | DAPI500-1 | volume | 820132220.00 |
|  | DAPI500-2 | volume | 1014072960.00 |
|  |  | sum | 1834205180.00 |
|  | DAPI-450 | volume | 741078660.00 |
|  | DAPI-350 | volume | 730276290.00 |
|  | DAPI-200 | volume | 362773312.00 |
|  |  | sum | 1834128262.00 |

**Supplemental Table 8.**
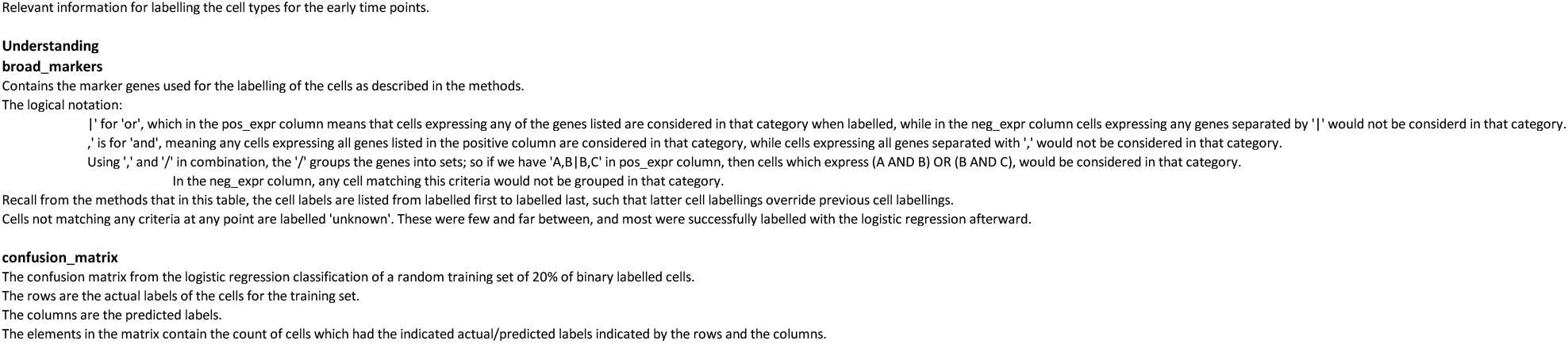
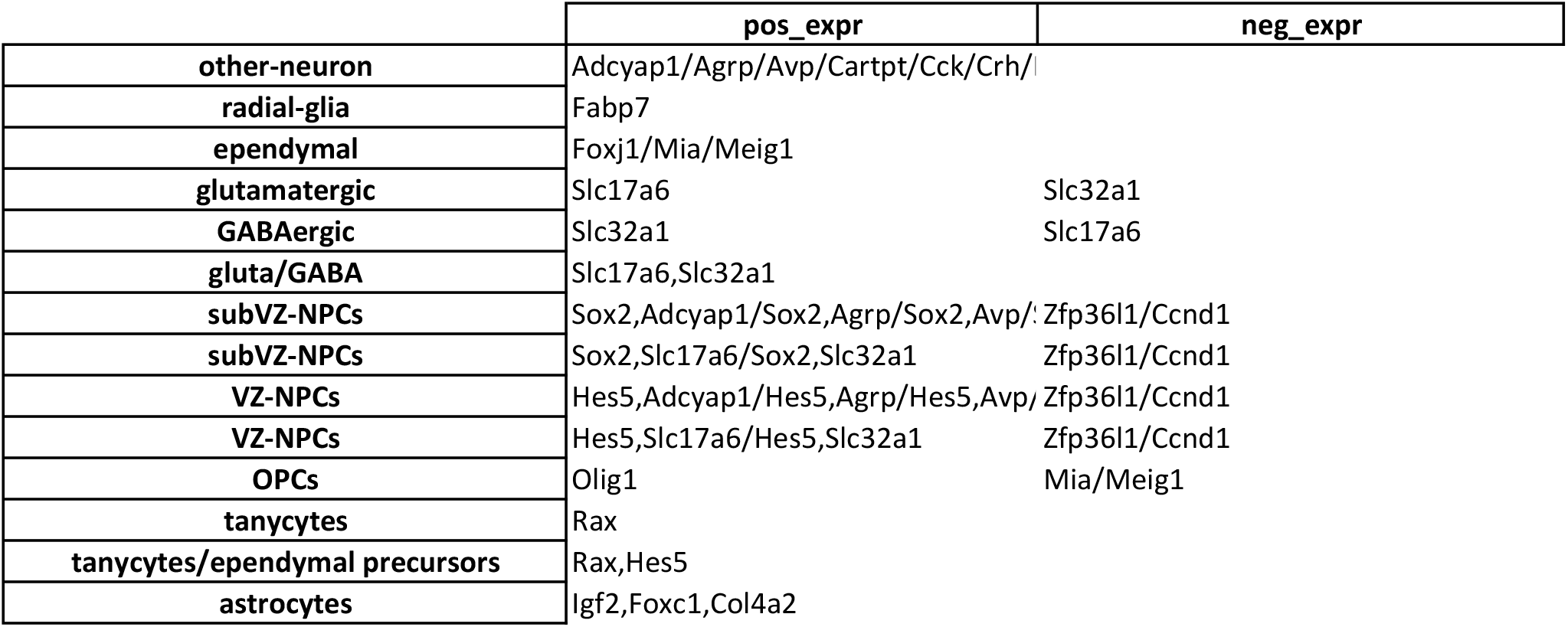
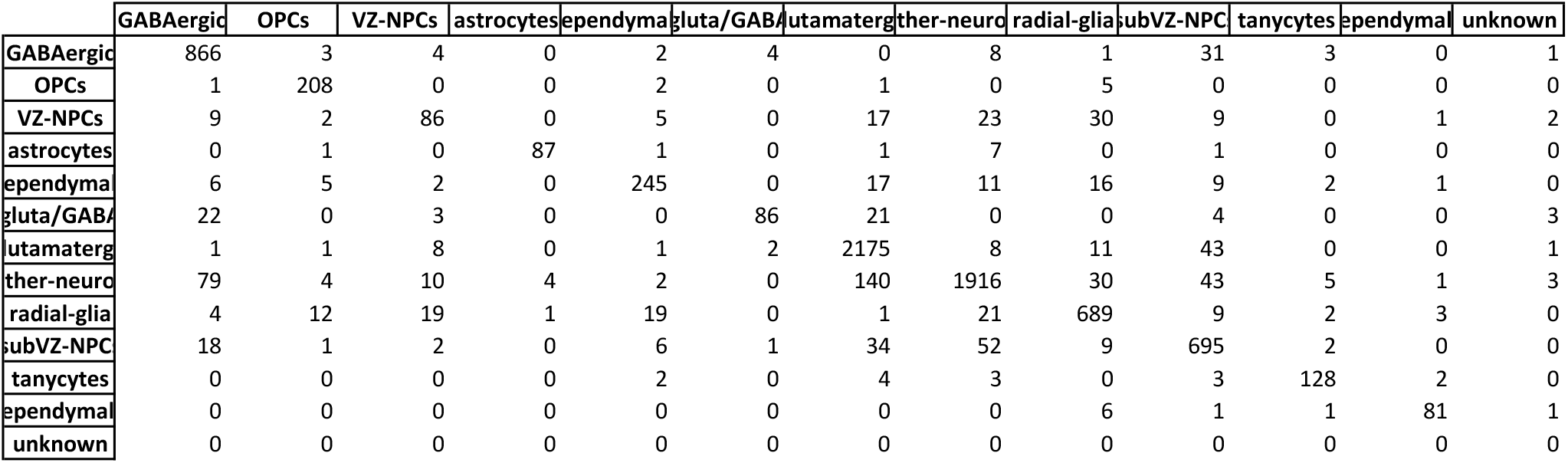
Marker genes used for broad labelling of early time points and confusion matrix from logistic regression classification. Relevant information for labelling the cell types for the early time points.

**Supplemental Table 9.** Differential expression between *Eed-cKO* and control in Lhx9 and Glut/GABA cells. Differentially expressed gene lists and associated statistics from comparing selected subsets of *Eed-cKO* cells against control from E13.5, E15.5, and E18.5. Lhx9 *Eed-cKO* cells were compared against Lhx9 control cells, and Glut/GABA *Eed-cKO* cells were compared against Glut/GABA control cells.

|  | **tvals** | **pvals** | **pvals_adj** | **logFC** |
| --- | --- | --- | --- | --- |
| **Hoxa9** | 33.33638 | 2.53E-18 | 1.21E-15 | 30.75007 |
| **Hoxa5** | 32.75009 | 3.52E-18 | 1.38E-15 | 29.38509 |
| **Meis2** | 32.31477 | 1.19E-25 | 1.51E-21 | 3.673903 |
| **Tal1** | 31.06243 | 5.58E-18 | 2.09E-15 | 7.476678 |
| **Hoxb7** | 29.79492 | 2.06E-17 | 6.36E-15 | 29.00135 |
| **Hoxc8** | 29.17854 | 3.03E-17 | 8.92E-15 | 28.77946 |
| **Bmi1** | 26.31524 | 1.72E-25 | 1.51E-21 | 2.119488 |
| **Hddc3** | 26.14458 | 2.49E-24 | 1.46E-20 | 3.086747 |
| **Meis1** | 25.84651 | 2.53E-22 | 4.47E-19 | 3.487031 |
| **Hoxa10** | 25.15477 | 4.59E-16 | 9.31E-14 | 10.48788 |
| **Hoxc9** | 24.22229 | 9.54E-16 | 1.81E-13 | 29.19356 |
| **Onecut2** | 24.21285 | 2.32E-19 | 1.57E-16 | 2.561906 |
| **Tfap2b** | 23.99061 | 4.49E-19 | 2.93E-16 | 4.887017 |
| **Hoxc13** | 23.76941 | 1.35E-15 | 2.47E-13 | 29.3826 |
| **Vax2** | 23.39516 | 1.28E-15 | 2.39E-13 | 7.895493 |
| **Crabp1** | 23.01755 | 3.87E-16 | 8.03E-14 | 5.718598 |
| **Tpt1** | 22.91051 | 9.98E-24 | 4.4E-20 | 1.372062 |

**Supplemental Table 10.** Differential expression between *Eed-cKO* Lhx9 and Glut/GABA cells against all other *Eed-cKO* cells. Differentially expressed gene lists and associated statistics from comparing selected subsets of *Eed-cKO* cells against the all other *Eed-cKO* cells from E13.5, E15.5, and E18.5. Lhx9 *Eed-cKO* cells were compared against all other *Eed-cKO* cells (excluding Glut/GABA *Eed-cKO*), and Glut/GABA *Eed-cKO* cells were compared against all other *Eed-cKO* cells (excluding Lhx9 *Eed-cKO*).

|  | **tvals** | **pvals** | **pvals_adj** | **logFC** |
| --- | --- | --- | --- | --- |
| **Lhx9** | 117.5029 | 1.17E-28 | 1.03E-24 | 32.75817 |
| **Nr2f1** | 18.25376 | 3.52E-20 | 1.55E-16 | 2.057193 |
| **Ebf1** | 16.97099 | 2.03E-18 | 5.98E-15 | 2.465343 |
| **Tcf7l2** | 16.15972 | 1.68E-15 | 2.7E-12 | 2.280319 |
| **Ebf3** | 16.08967 | 4.84E-18 | 1.07E-14 | 2.429213 |
| **Vim** | 15.97195 | 2.71E-18 | 6.82E-15 | 2.06231 |
| **Rpl7** | 14.37115 | 5.41E-15 | 6.36E-12 | 0.945302 |
| **Nhlh2** | 13.89431 | 2.31E-15 | 3.13E-12 | 2.147537 |
| **Barhl2** | 13.36624 | 7.58E-16 | 1.34E-12 | 2.107711 |
| **Lhx2** | 13.34695 | 3.47E-14 | 3.06E-11 | 1.813185 |
| **Tpt1** | 12.8143 | 8.18E-14 | 6.87E-11 | 1.247123 |
| **Ube2c** | 12.6677 | 8.12E-15 | 8.43E-12 | 3.084001 |
| **Eef1a1** | 12.51484 | 5.14E-13 | 2.92E-10 | 0.735164 |
| **Cnn3** | 12.31295 | 2.22E-13 | 1.53E-10 | 1.492037 |
| **Hbb-y** | 12.02595 | 6.73E-13 | 3.71E-10 | 2.678972 |
| **Hes5** | 11.98529 | 2.35E-14 | 2.3E-11 | 2.46236 |
| **Nde1** | 11.89173 | 1.99E-13 | 1.53E-10 | 3.535015 |
| **Cenpa** | 11.33605 | 2.39E-12 | 1.03E-09 | 2.477243 |
| **Crabp2** | 11.30777 | 1.06E-12 | 5.33E-10 | 2.162725 |
| **H1f0** | 11.29994 | 2.84E-12 | 1.19E-09 | 1.297954 |
| **Ccna2** | 11.19255 | 3.76E-13 | 2.29E-10 | 2.551879 |
| **Tcf12** | 10.87684 | 5.83E-12 | 2.06E-09 | 1.267464 |
| **Ddah2** | 10.81243 | 6.38E-12 | 2.16E-09 | 1.406031 |
| **Rps9** | 10.79055 | 3.73E-11 | 1.03E-08 | 0.941671 |
| **Rps11** | 10.77306 | 1.56E-11 | 4.58E-09 | 0.989904 |
| **Sdcbp** | 10.61393 | 2.32E-12 | 1.02E-09 | 1.072293 |
| **Hmgb2** | 10.567 | 7.26E-13 | 3.88E-10 | 2.281737 |

**Supplemental Table 11.**
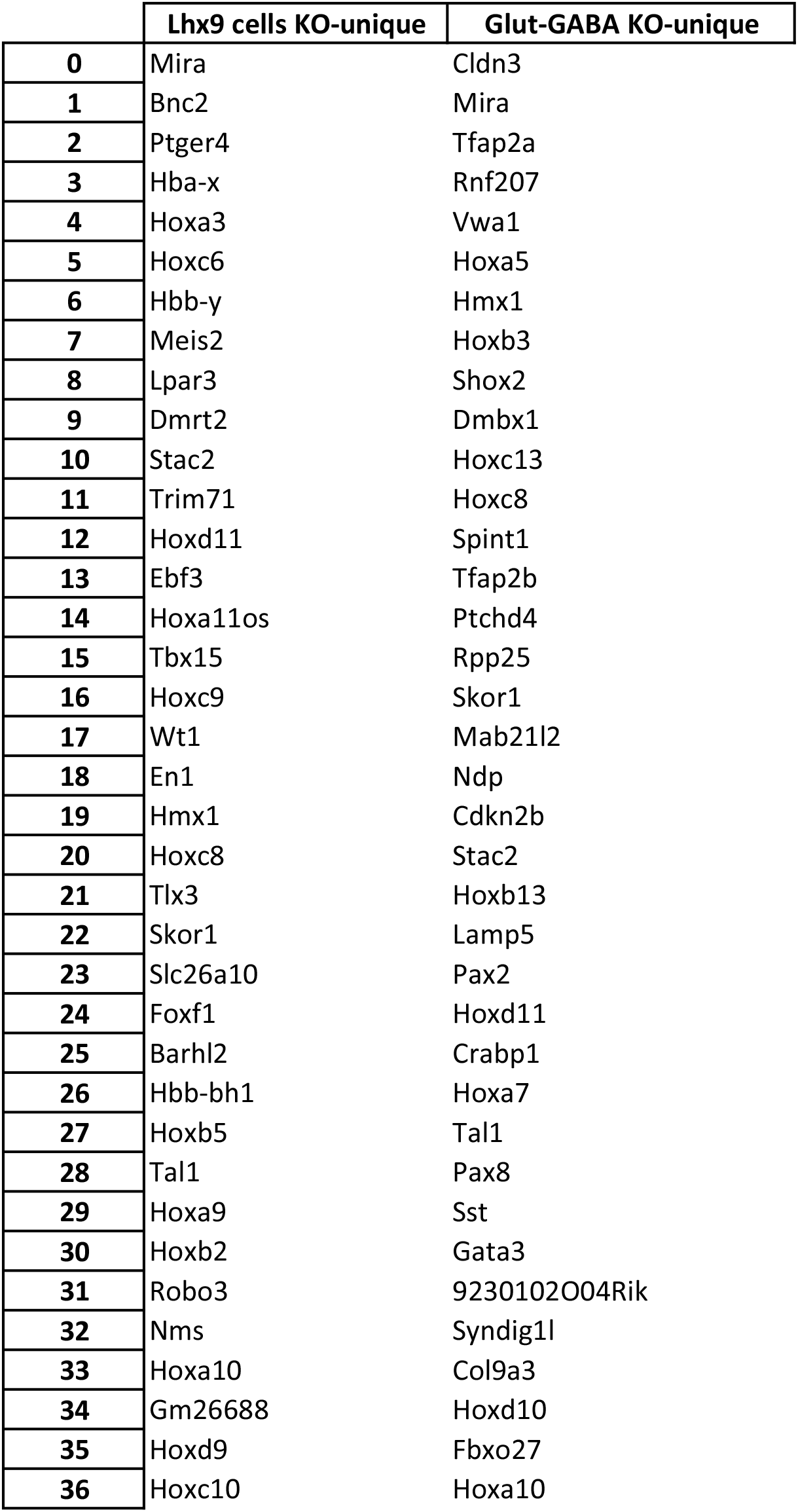
Genes differentially effected in Lhx9 and Glut/GABA *Eed-cKO* cells but not other cells. Overlap of the gene lists from comparing Lhx9 *Eed-cKO* versus control and Lhx9 *Eed-cKO* cells versus all other *Eed-cKO* cells. Also overlap genes from comparing Glut/GABA *Eed-cKO* versus control and Glut/GABA *Eed-cKO* cells versus all other *Eed-cKO* cells. These genes thereby represent those which were significantly *Eed-cKO* effected in these cell subsets but not all other cells.

